# A rapid and bidirectional reporter of neural activity reveals neural correlates of social behaviors in *Drosophila*

**DOI:** 10.1101/2023.04.10.536242

**Authors:** Moise Bonheur, Kurtis J. Swartz, Melissa G. Metcalf, Anna Zhukovskaya, Avirut Mehta, Kristin E. Connors, Julia G. Barasch, Xinke Wen, Andrew R. Jamieson, Kelsey C. Martin, Richard Axel, Daisuke Hattori

## Abstract

Neural activity is modulated over different timescales encompassing sub-seconds to days reflecting changes in external environment, internal state, and behavior. Using *Drosophila* as a model, we have developed a rapid and bidirectional reporter that provides a robust cellular readout of recent neural activity. This reporter utilizes nuclear vs cytoplasmic distribution of CREB-regulated transcriptional coactivator, CRTC. Subcellular distribution of GFP-tagged CRTC (CRTC::GFP) bidirectionally changes on the order of minutes and reflects both increases and decreases in neural activity. We establish an automated machine-learning-based routine for efficient quantification of reporter signal. Using this reporter, we demonstrate acute mating- evoked activation of peptidergic neurons. We further investigate the functional role of the master courtship regulator gene, *fruitless*, and show that *fruitless* is necessary to ensure activation of male arousal neurons by female cues. Together, our results establish CRTC::GFP as a bidirectional reporter of recent neural activity suitable for examining neural correlates in behavioral contexts.

## INTRODUCTION

Neural activity is optimally examined by electrophysiological or optical recording, which resolves neural activity on a fast timescale of sub-seconds. These recording methods require tethering each animal to a recording device and therefore can hinder the manifestation of natural behaviors. In addition, the throughput of these methods is typically low. Transcriptional reporters of neural activity provide an efficient method to identify neurons that are activated during natural behaviors ^1^. These reporters utilize activity-dependent gene expression to label active neurons. Some reporters are endogenous proteins or transcripts encoded by immediate early genes ^2–4^.

Engineered reporters include those that use the enhancer of immediate early genes to express marker proteins ^5–8^ and those that employ exogenous transcription factors whose activation is regulated by neural activity ^9, 10^. Because these reporters rely on detection of gene expression, they operate on the timescale of hours ^4, 9, 10^. This timescale obscures the temporal correlation between detected neural activation and its causal event. Moreover, the stability of reporter proteins or transcripts is unrelated to neural activity and therefore these reporters remain in neurons long after causal neural activity terminates.

We have developed a bidirectional neural activity reporter that operates on the timescale of minutes using the *Drosophila* model. This reporter takes advantage of activity-dependent translocation of CRTC (CREB-regulated transcriptional coactivator) from the cytoplasm to the nucleus. In both insect and mammalian cells, phosphorylated CRTC is sequestered in the cytoplasm through binding to a scaffold protein, 14-3-3 ^11–15^. Dephosphorylated CRTC translocates into the nucleus to regulate CREB-mediated transcription. Phosphorylation of CRTC is regulated by salt-inducible kinase 2 (SIK2) and calcineurin phosphatase. CRTC activation is mediated by cAMP signaling, which inhibits SIK2 kinase activity, and by calcium signaling, which activates calcineurin. In mouse primary neuronal cultures and hippocampal slices, CRTC1 translocates into the nucleus upon synaptic activation or electrical stimulation that induces long-term potentiation ^16, 17^, acting as a cytoplasm-to-nucleus messenger that contributes to activity-dependent induction of CREB target genes.

In this study, we determine that fly CRTC translocates into the nucleus upon increased neural activity and out of the nucleus upon diminished neural activity on the order of minutes and demonstrate its utility as a rapid and bidirectional reporter of neural activity. Using *in vivo* 2- photon imaging, we first show that GFP-tagged CRTC (CRTC::GFP) translocates from the cytoplasm into the nucleus following neuronal excitation induced by olfactory stimuli. We develop an automated quantification method using deep convolutional neural networks. We then use CRTC::GFP to investigate neural activity in two behavioral contexts, mating and courtship. Finally, we provide evidence that CRTC::GFP can be used to study neuronal activity in the context of starvation. Together, our results establish CRTC::GFP as a robust bidirectional reporter of recent neural activity that is broadly useful for examining neural correlates of external sensory environment, internal state, and behavioral state.

## RESULTS

### *Drosophila* CRTC translocates into the nucleus upon neural activation

We first examined whether neuronal activation drives nuclear translocation of fly CRTC *in vivo* (Fig. 1a). We generated a GFP-tagged CRTC construct (CRTC::GFP) and, using the GAL4/UAS system ^18^, expressed this fusion protein in a class of second-order olfactory projection neurons (PNs) that extend their dendrites into the DC3 glomerulus of the antennal lobe. The olfactory sensory neurons that innervate this glomerulus robustly respond to farnesol, an odor found in citrus fruit peels ^19^. We confirmed that DC3 PNs are activated by farnesol using *in vivo* 2-photon calcium imaging (Extended Data Fig. 1a) with a genetically encoded calcium indicator, GCaMP6f ^20^. We then utilized the same *in vivo* 2-photon imaging setup and, instead of GCaMP signals, monitored the subcellular distribution of CRTC::GFP signals in the cell bodies of DC3 PNs. CRTC::GFP was expressed in these neurons using a split-GAL4 (spGAL4, ^21^) driver, SS01165 ^22^. A membrane-targeted red fluorescent protein, mCD8::mCherry (mouse CD8 fused to mCherry), was coexpressed to demarcate plasma and nuclear membranes such that subcellular distribution of CRTC::GFP signals could be quantified. CRTC::GFP signals in the nucleus strongly increased after farnesol presentations (Fig. 1b), indicating that activation of DC3 neurons results in nuclear translocation of CRTC molecules.

**Fig. 1:**
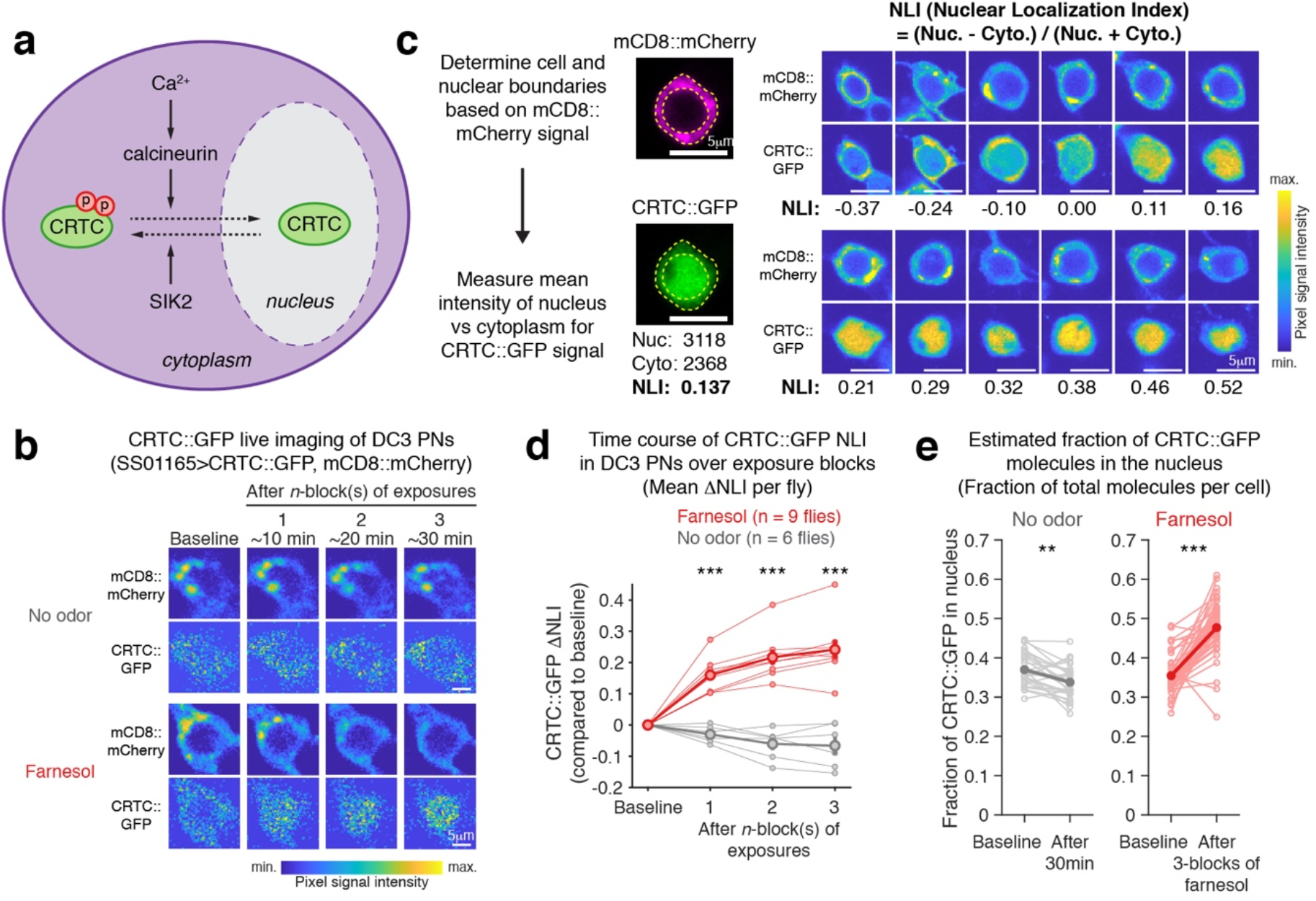
*Drosophila* CRTC translocates into the nucleus upon odor-evoked neural activation. (a) Schematic representation of CRTC translocation. In a basal state, CRTC is phosphorylated by salt-induced kinase (SIK2) and localizes in the cytoplasm. A calcium-dependent phosphatase, calcineurin, dephosphorylates CRTC, resulting in nuclear translocation of CRTC. (b) Representative 2-photon images of mCD8::mCherry and CRTC::GFP in DC3 cell bodies. Flies were exposed to three blocks of farnesol presentations (see Methods). Including imaging time, each block was approximately 10 min. Images for control condition with no odor were taken every 10 min. See Supplementary Table 1 for genotypes. (c) Quantification of nuclear localization index (NLI). *Left*; mCD8::mCherry is used to demarcate plasma and nuclear membranes in the cell body (yellow dotted lines). Mean pixel GFP signal intensity is calculated in the regions-of-interest for nucleus and cytoplasm. NLI is calculated by (mean nuclear GFP signal – mean cytoplasmic GFP signal) / (mean nuclear GFP signal + mean cytoplasmic GFP signal) and ranges from -1 to +1. *Right*; example images of cell bodies that exhibit different CRTC localization patterns with corresponding NLIs. Example cell images are single confocal slices of immunostained cell bodies (see Methods). (d) Changes in NLI (τιNLI) of DC3 PNs upon odor exposure. Thin lines; individual flies, thick lines; mean ± SEM. See Extended Data Fig. 1b for raw NLIs for individual flies. Statistics by Wilcoxon rank sum test; *** p<0.001. Statistics including exact p-values are in Supplementary Table 2. (e) Estimated fractions of CRTC::GFP molecules in the nucleus per cell body. Molecular fractions are calculated based on NLIs observed for baseline and after three blocks of farnesol presentations with relative nuclear radius of 0.7. See Extended Data Fig. 1c and Methods. N = 29 cells (no odor) and 48 cells (farnesol). Statistics by Wilcoxon signed rank test; ** p<0.01, *** p<0.001.

We quantified subcellular distribution of CRTC::GFP in each neuron by calculating the nuclear localization index (NLI) using the formula, (mean nuclear GFP signal – mean cytoplasmic GFP signal) / (mean nuclear GFP signal + mean cytoplasmic GFP signal). The NLI ranges from -1 to +1 and indicates the relative distribution of CRTC::GFP inside and outside the nucleus in individual neurons (Fig. 1c). The NLI strongly increased after the first block of farnesol presentations (difference in NLI between pre-stimulus baseline and post-stimulus (ι1NLI) for farnesol 0.16±0.05 vs no-odor control -0.03±0.03, p<0.001, Mean ± SD, Fig. 1d and Extended Data Fig. 1b, see Methods and Supplementary Table 2 for statistics). The NLI further increased upon exposure to subsequent blocks of farnesol presentations (ι1NLI after three blocks of farnesol 0.24±0.09, p<0.05 between farnesol exposure blocks). By contrast, the NLI did not increase in control flies that were not presented with farnesol for the duration of experiments. These results demonstrate that CRTC::GFP translocates into the nucleus following neuronal activation.

We estimated the fraction of CRTC::GFP molecules that translocate into the nucleus following DC3 activation. We first determined the average radius of DC3 nuclei relative to that of cell bodies using the images acquired in our live imaging experiments and derived the ratio of nuclear vs cytoplasmic volumes (see Methods). We then calculated the fraction of CRTC::GFP molecules in the nucleus vs cytoplasm with an assumption that mean CRTC::GFP signal intensities correlate with concentrations of CRTC::GFP molecules in each subcellular compartment (Fig. 1e and Extended Data Fig. 1c). We found that on average 35.5±4.8% of total CRTC::GFP molecules were in the nucleus of DC3 neurons in the unstimulated baseline state. Upon farnesol presentations, this fraction increased to 47.7±6.8%. Thus, these calculations indicate that farnesol-induced activation of DC3 neurons resulted in nuclear translocation of approximately 19% of cytoplasmic CRTC::GFP molecules. Together, we conclude that neural activation drives nuclear translocation of fly CRTC.

### Subcellular distribution of CRTC::GFP provides a readout of neural activity in free moving flies

Our *in vivo* 2-photon imaging experiments showed activity-dependent nuclear translocation of CRTC::GFP, but these experiments required tethering the flies and were low-throughput. We therefore employed immunostaining and confocal microscopy, a method that provides greater signal-to-noise ratio as well as higher throughput, and tested the utility of CRTC::GFP in reporting neural activity in free moving flies (see Methods). We first examined whether exogenous neural activation using a heat-gated cation channel, dTRPA1 ^23^, induces nuclear localization of CRTC::GFP. Two different classes of neurons were independently examined. The first, PAM-γ4, is a class of dopaminergic neurons that innervates the mushroom body, an olfactory center of the fly. We expressed CRTC::GFP and mCD8::mCherry together with dTRPA1 in PAM-γ4 neurons using the MB312B spGAL4 driver ^24^. We then activated these neurons by incubating flies at 32°C for one hour and immunostained the brains of these flies to visualize the subcellular distribution of CRTC::GFP signals by confocal microscopy (Fig. 2a).

**Fig. 2:**
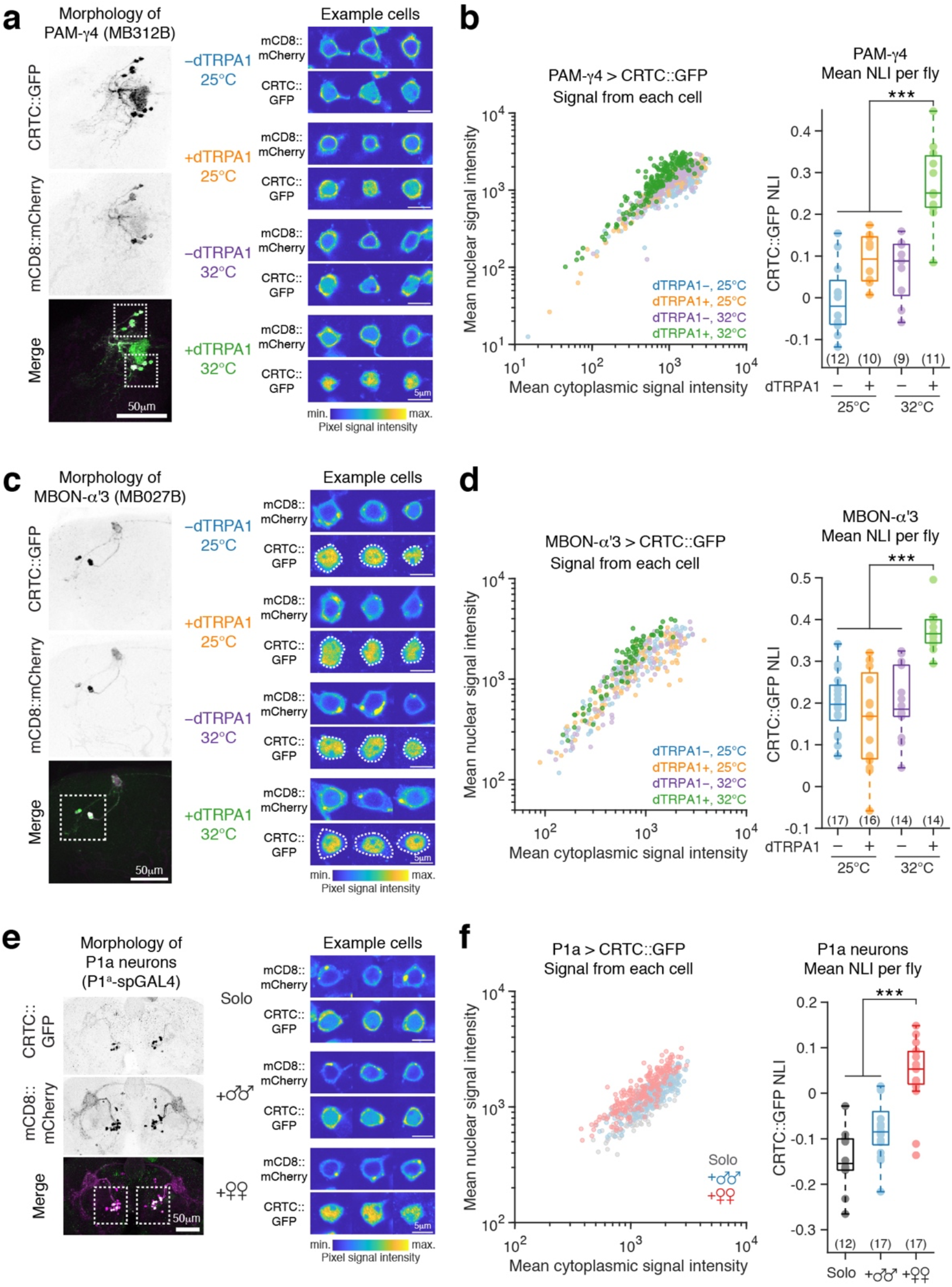
Subcellular distribution of CRTC::GFP provides a readout for neural activity in free moving flies. (a) Example confocal images of PAM-γ4 from thermogenetic activation experiments. *Left*; morphology of PAM-γ4. Dotted areas contain cell bodies. In merged image, green; CRTC::GFP, magenta; mCD8::mCherry. *Right*; representative example images of single PAM-γ4 cell bodies in different conditions. Each image is a confocal slice and mCD8::mCherry (top) and CRTC::GFP (bottom) signals are shown. (b) Thermogenetic activation experiments for PAM-γ4. Experimental flies with or without dTRPA1 were incubated at 25°C or 32°C for one hour before brain dissection, fixation, and immunostaining. *Left*; mean cytoplasmic vs nuclear CRTC::GFP signal intensities for each cell. *Right*; mean CRTC::GFP NLI averaged across cells per fly. Each dot represents a fly (number of flies in parenthesis) and boxplot shows median (horizontal line), first and third quartiles (box), and data within 1.5x IQR (whiskers). See Extended Data Fig. 2a for individual fly data. Kruskal- Wallis test followed by a post hoc Tukey’s HSD test; *** p<0.001. (c) *Left*; morphology of MBON-α’3 with dotted area indicating cell bodies. *Right*; representative confocal slice images of MBON-α’3 from thermogenetic activation experiments. Dotted outlines in CRTC::GFP images indicate plasma membrane of each cell determined in mCD8::mCherry images. (d) Thermogenetic activation experiments for MBON-α’3. Experimental flies were incubated at 25°C or 32°C for one hour before brain dissection, fixation, and immunostaining. *Left*; mean cytoplasmic vs nuclear CRTC::GFP signal intensities for each cell. *Right*; mean CRTC::GFP NLI averaged across cells per fly. See Extended Data Fig. 2d for individual fly data and Extended Data Fig. 2e for GFP only controls. Kruskal-Wallis test followed by a post hoc Tukey’s HSD test; *** p<0.001. (e) Example confocal images of P1a neurons. Subject flies were paired for three days with males or females, or were left single-housed (“solo”). *Left*; morphology of P1a neurons with dotted areas indicating cell bodies. *Right*; representative confocal slice images of single P1a cell bodies in different conditions. (f) Effect of different rearing conditions upon CRTC::GFP localization in P1a neurons. *Left*; mean cytoplasmic vs nuclear CRTC::GFP signal intensities for each cell. *Right*; mean CRTC::GFP NLI averaged across cells per fly. See Extended Data Fig. 2f for individual fly data and Extended Data Fig. 2g for data obtained with one day of social rearing conditions. Statistics by Kruskal-Wallis test followed by a post hoc Tukey’s HSD test; *** p<0.001.

Our analyses revealed that thermogenetically activated PAM neurons exhibited higher nuclear CRTC::GFP signals (NLI 0.27±0.10) than PAM neurons in control flies that were not exposed to heat (NLI 0.09±0.06) or lacked the UAS-dTRPA1 transgene (NLI 0.07±0.08, or both NLI - 0.01±0.09, p<0.001, Fig. 2b and Extended Data Fig. 2a).

We also studied a class of mushroom body output neurons, MBON-α’3, labeled by the MB027B spGAL4 driver ^24^. Under control conditions, these neurons exhibited higher basal nuclear localization of CRTC::GFP than PAM-γ4 neurons (Fig. 2a vs c). RNA interference experiments indicate that the basal nuclear localization in MBON-α’3 is in part mediated by calcineurin phosphatases (Extended Data Fig 2b-c, see below). Despite the high basal nuclear localization, thermogenetically activated MBON-α’3 exhibited even higher nuclear CRTC::GFP signals than those in control conditions (NLI for dTRPA1+(32°C) 0.37±0.05, dTRPA1+(25°C) 0.16±0.12, dTRPA1–(32°C) 0.20±0.08, dTRPA1– (25°C) 0.20±0.07, p<0.001, Fig. 2c-d and Extended Data Fig. 2d-e). These observations indicate that neural activation can be detected using CRTC::GFP even in the presence of high basal nuclear localization.

We next tested the sensitivity of CRTC::GFP in detecting differences in physiological levels of neural activity. We employed neurons found in the P1 cluster of the male courtship circuit as a model. Neurons in the P1 cluster are male-specific and play an important role in mediating social behaviors of male flies, such as female-directed courtship and male-directed aggression ^25–34^. P1 neurons are activated by courtship-promoting chemosensory and visual stimuli, such as contact-dependent female pheromones and fly-sized moving visual cues ^27, 31, 35–37^. Thus, P1 neuron activity is thought to represent the state of arousal in males typically induced by the presence of females. We expressed CRTC::GFP together with mCD8::mCherry in a subset of P1 neurons (P1a neurons) using a spGAL4 driver, P1^a^-spGAL4 ^29, 30^. These experimental male flies were housed for three days either singly, grouped with male partners, or grouped with female partners. We observed greater nuclear CRTC::GFP signals in males housed with female partners compared to males housed singly or with male partners (NLI for female-paired flies 0.04±0.08, single-housed flies -0.15±0.07 and for male-paired flies -0.08 ±0.06, p<0.001, Fig. 2e-f and Extended Data Fig. 2f). Higher CRTC::GFP nuclear signals were also observed after one day of cohousing with female partners (NLI for female-paired flies 0.07±0.08, single-housed flies -0.12±0.11, and male-paired flies -0.11±0.05, p<0.05, Extended Data Fig. 2g). These observations indicate that the presence of females in the environment results in the activation of P1a neurons and translocation of CRTC into the nucleus. The subcellular distribution of CRTC::GFP therefore provides a sensitive measure of neural activity in free moving flies under physiological conditions.

### CRTC::GFP redistributes on the order of minutes and indicates both increases and decreases in neural activity

We examined the kinetics of nuclear import using thermogenetic activation of MBON-α’3. Five minutes of thermogenetic activation was sufficient to observe a significant increase in nuclear CRTC::GFP signals as assessed with immunostaining and confocal imaging (NLI for 5-minute activation 0.32±0.07 vs no-activation 0.16±0.10, p<0.001, Fig. 3a). Elevated nuclear signals were also observed after two hours of continued heat exposure (NLI after 2-hour 32°C incubation for dTRPA1-positive flies 0.34±0.07, for dTRPA1-negative flies 0.14±0.12, p<0.05). We next examined the kinetics of nuclear export by thermogenetically activating MBON-α’3 for 15 minutes and then transferring the flies to room temperature (RT, 22-25°C). Nuclear CRTC::GFP signals rapidly diminished over time and were indistinguishable from the baseline level after 30-minute (NLI for baseline 0.19±0.10, right after 15-minute 32°C incubation 0.38±0.06 and after 30-minute RT incubation 0.19±0.09, Fig. 3b). Together, our time course experiments demonstrate that the nuclear localization of CRTC::GFP increases within minutes upon neural activation, persists over hours under continual activation, and returns to a baseline within 30 minutes following termination of neural activity.

**Fig. 3:**
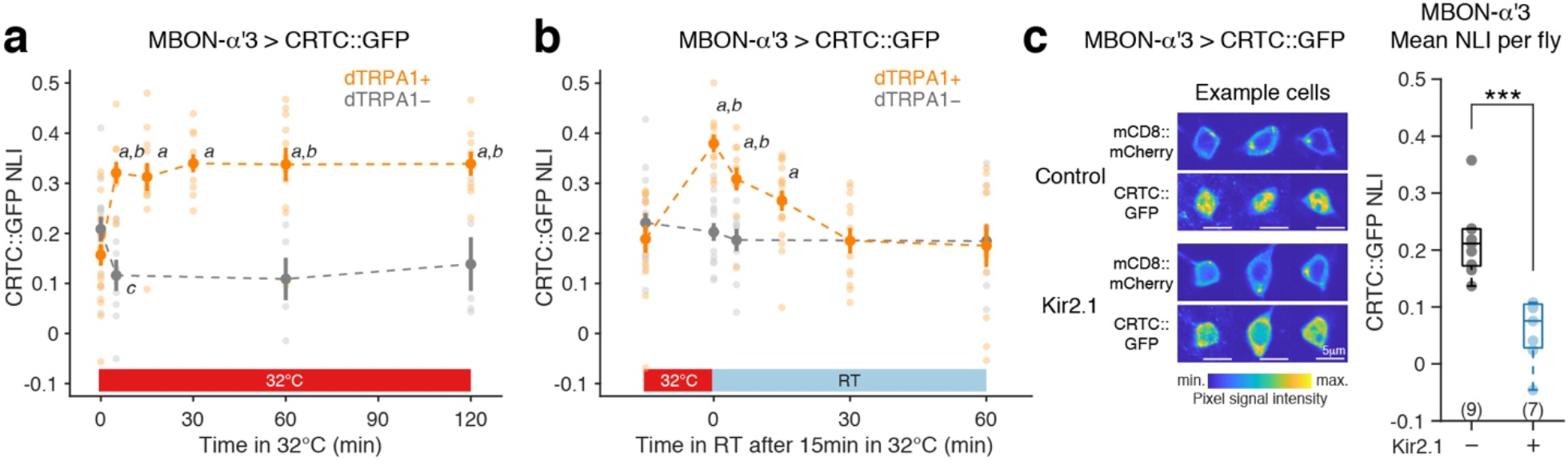
Subcellular distribution of CRTC::GFP changes on the order of minutes and indicates both increases and decreases in neural activity. (a) Time-course experiments that examine nuclear import kinetics of CRTC::GFP. Flies with or without dTRPA1 expression in the MBON-α’3 were incubated at 32°C for 0, 5, 15, 30, 60, or 120 min before dissection, fixation, and immunostaining. Statistics by Wilcoxon rank sum test; *a* denotes p<0.05 compared to dTRPA1+ baseline (0 min incubation), *b* denotes p<0.05 compared to dTRPA1- at each timepoint, and *c* denotes p=0.035 compared to dTRPA1- baseline (0 min incubation). N=9-20 for all timepoints except dTRPA1- 60 and 120 min, for which N=5 each. (b) Time-course experiments that examine nuclear export kinetics of CRTC::GFP. Flies with or without dTRPA1 expression in MBON-α’3 were incubated at 32°C for 15 min then were incubated at room temperature (22-25°C) for 0, 5, 15, 30, or 60 min before dissection, fixation, and immunostaining. Statistics by Wilcoxon rank sum test; *a* denotes p<0.05 compared to dTRPA1+ baseline (no incubation at 32°C), and *b* denotes p<0.05 compared to dTRPA1- at each time point. N=11-19. (c) Genetic silencing experiments of MBON-α’3. MBON-α’3 was silenced by overexpression of Kir2.1 using the MB027B driver. *Left*; representative confocal images. *Right*; each dot represents mean NLI across cells per fly (number of flies in parenthesis). Statistics by Wilcoxon rank sum test; *** p<0.001.

Because neural activity is represented by the relative distribution of CRTC::GFP signals in the nucleus vs the cytoplasm, this may allow for detecting a decrease in neural activity when neurons are inhibited or silenced. We tested this possibility by examining CRTC::GFP signals in MBON-α’3 that were silenced by overexpression of an inward-rectifying potassium channel, Kir2.1 ^38^. We generated an untagged version of the UAS-Kir2.1 transgene and expressed it together with CRTC::GFP and mCD8::mCherry in MBON-α’3 using the MB027B driver. We observed significantly lower nuclear CRTC::GFP signals in silenced MBON-α’3 as compared to MBONs in control flies that did not express Kir2.1 (NLI for Kir2.1 flies 0.06±0.06 vs controls 0.22±0.06, p<0.001, Fig. 3c). Thus, these results demonstrate that CRTC::GFP localizes to the cytoplasm in neurons that are silenced, providing bidirectionality in reporting neural activity.

Together with the observation that calcineurins contribute to determining basal levels of CRTC::GFP in the nucleus (Extended Data Fig. 2b-c), these results also suggest that the baseline nuclear localization of CRTC::GFP is determined by basal calcium levels that may reflect tonic neural activity.

In summary, our results establish CRTC::GFP as a subcellular-distribution-based bidirectional neural activity reporter that operates on the timescale of minutes, which can be used to detect and monitor both increases and decreases in neural activity.

### Automating quantification of CRTC::GFP localization using machine learning

A bottleneck for efficient use of the CRTC::GFP reporter is determination of the nuclear vs cytoplasmic regions-of-interest (ROIs) to perform signal quantification, which requires manually drawing cellular and nuclear boundaries based on mCD8::mCherry signals (Fig. 1c). We improved the utility of this reporter by automating the subcellular ROI determination.

Because the intensity and distribution of the mCD8::mCherry marker varies between neurons and between flies (for example, see Fig. 4a and c), simple image processing methods such as thresholding are not suitable. We therefore employed machine-learning-based image segmentation. Labeling the nucleus and the cytoplasm in two different colors provides additional signals to more precisely determine subcellular ROIs in each neuron. We therefore generated a bicistronic transgene (UAS-mCD8::mCherry-T2A-nls::LacZ) expressing nuclear-localized ý- galactosidase (nls::LacZ, nls - nuclear localization signal) and mCD8::mCherry. We then expressed this transgene together with CRTC::GFP in 16 neuron types that encompass different shapes and sizes and manually determined the cellular and nuclear boundaries using mCD8::mCherry and nls::LacZ signals for over 9,000 neurons. This generated a dataset appropriate for establishing machine-learning-based image segmentation.

**Fig. 4:**
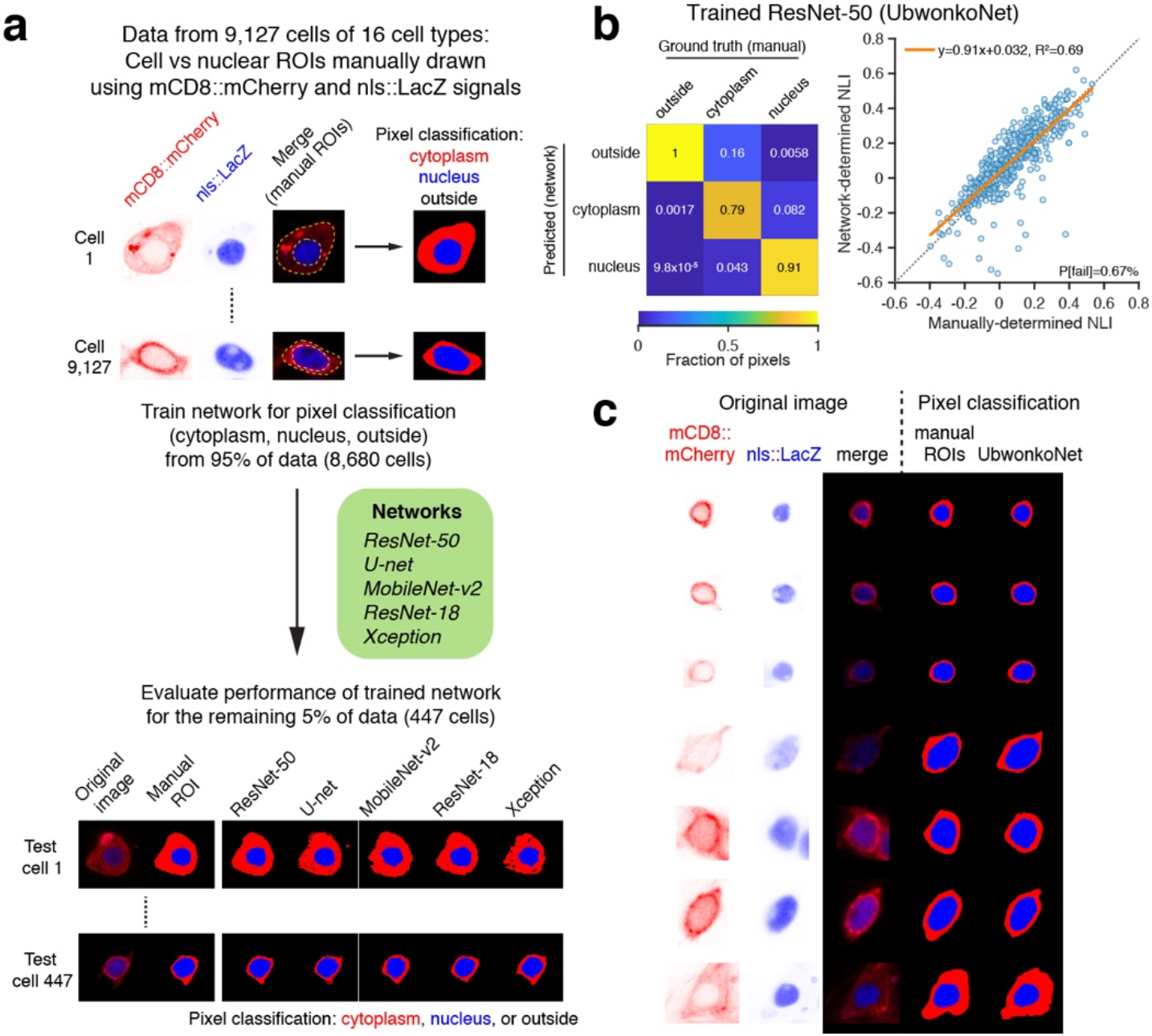
Automating pixel classification by deep convolutional neural network. (a) Workflow for training different convolutional neural networks and evaluation of performance. Membrane (mCD8::mCherry) and nuclear (nls::LacZ) markers were expressed from a bicistronic transgene, UAS-mCD8::mCherry-T2A-nls::LacZ. Confocal slice images comprising membrane and nuclear signals (i.e., no CRTC::GFP signal) and their pixel classification based on manually-drawn ROIs were the input for training each network. We trained 5 different networks using 95% (8,680) of cells and the performance of each network was evaluated for 5% (447) of held-out cells. Pixel classification for two example test cells are shown at the bottom. (b) Performance evaluation for trained ResNet-50. See Extended Data Fig. 3a for the other networks. *Left*; confusion matrix for pixel-level performance evaluation. *Right*; comparison of NLIs for CRTC::GFP signal determined manually vs by network. Each dot is a cell and the orange line is regression. P[fail] denotes percentage of cells that each network fails to assign both cytoplasm and nucleus. See also Supplementary Table 3 for other metrics of network performance. (c) Seven examples of pixel classification by trained ResNet-50 (UbwonkoNet). See Extended Data Fig. 3b for additional examples.

We trained five different convolutional neural networks to classify pixels in the images of neuronal cell bodies as nuclear, cytoplasmic, or outside the cell (Fig. 4a) and assessed the performance of each trained network in two different ways; first at the pixel level by confusion matrix and second at the level of NLI (Fig. 4b and Extended Data Fig. 3a, see also Supplementary Table 3 for more metrics of network performance). The best performing network (“UbwonkoNet”) classified 79% of cytoplasmic pixels and 91% of nuclear pixels correctly.

Manually-determined vs network-determined NLIs closely correlated with each other (regression coefficient 0.91, intercept 0.03, coefficient of determination 0.69, Fig. 4b-c). By analyzing a dataset that examined female-dependent activation of P1a neurons, we confirmed that the same experimental result was obtained using either UbwonkoNet- or manually-determined NLIs (Extended Data Fig. 3c). Thus, we conclude that the UbwonkoNet can be used to automatically determine nuclear vs cytoplasmic ROIs, providing an efficient analysis method to utilize CRTC::GFP as a neural activity reporter.

### CRTC::GFP detects acute activation of peptidergic neurons upon mating

The rapid redistribution of CRTC::GFP may allow us to report and monitor the activity of neurons in behavioral contexts. Here, we focus on two social behaviors, mating and courtship, for which conventional recording methods may be inefficient or incompatible.

Interneurons in the abdominal ganglion of male flies that express the neuropeptide, corazonin (Crz), are necessary for male mating behavior ^39^. Male flies in which Crz neurons are genetically silenced are infertile and exhibit prolonged copulation, whereas activation of these neurons results in premature sperm transfer and shortened copulation. These results suggest that Crz neurons are activated during mating to regulate proper sperm transfer. We have tested this hypothesis by examining the subcellular distribution of CRTC::GFP in the Crz neurons of male flies that had just mated. Mating was monitored manually and, immediately after mating ended (copulation duration, 17.5±3.1 min, N=16), the abdominal ganglion of each male was dissected, fixed, and immunostained. Mated males exhibited higher nuclear CRTC::GFP signals in Crz neurons than those in control males that did not mate (NLI for mated males 0.01±0.09 vs unmated males -0.21±0.09, p<0.001, Fig. 5a). Expression of CRTC::GFP in Crz neurons did not significantly affect male mating behavior (Fig. 5b and Extended Data Fig. 4), suggesting that the expression of CRTC::GFP does not disrupt normal neuronal activity. These results demonstrate that Crz neurons are indeed activated by mating and highlight the utility of the CRTC::GFP reporter to monitor neural activity in behaving flies.

**Fig. 5:**
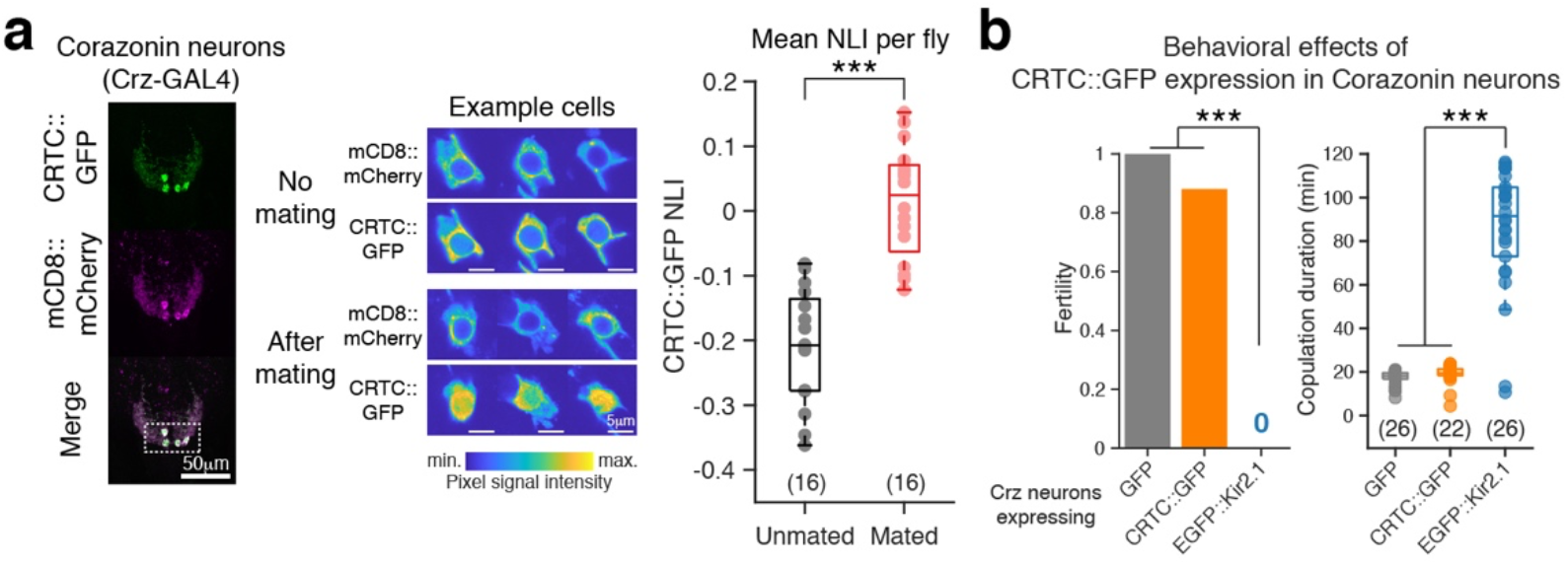
CRTC::GFP detects acute activation of peptidergic neurons upon mating. (a) Subcellular distribution of CRTC::GFP in corazonin neurons in male flies that had just mated vs not mated. *Left*; morphology of corazonin neurons in abdominal ganglion. Dotted area contains cell bodies. *Middle*; representative confocal slices of cell bodies. *Right*; NLI of CRTC::GFP signal in corazonin-expressing neurons of males that just mated vs unmated. Statistics by Wilcoxon rank sum text; *** p<0.001. (b) *Left*; fertility of flies expressing GFP, CRTC::GFP, or EGFP::Kir2.1 in corazonin neurons. N=25 each. Statistics by Fisher’s exact test and Bonferroni correction; *** p<0.001. *Right*; copulation duration in 2-hour observation period for fly pairs each consisting of a male of indicated genotype and a wild-type female. See Extended Data Fig. 4b for ethograms. Statistics by Kruskal-Wallis test followed by a post hoc Tukey’s HSD test; *** p<0.001.

### Examination of the activity of P1a neurons in *fruitless* mutants using CRTC::GFP

We next used CRTC::GFP to begin examining the neural basis underlying genetic specification of courtship behavior. The *Drosophila fruitless* (*fru*) gene is necessary for specifying female-directed courtship behavior displayed by male flies ^28, 40–45^. *fru* mutant males exhibit no courtship behavior toward females but do exhibit a subset of courtship rituals toward males ^46–49^. Little is known about neural basis of this behavioral transformation. Interestingly, P1 neurons, which mediate arousal in males and promote courtship, are present in *fru* mutants and optogenetic activation of P1 neurons in these mutants induces courtship behavior toward females as well as males ^34^. These observations suggest that altered courtship behavior in *fru* mutants may arise from altered activation of P1 neurons. We examined P1a activity in *fru* mutants using CRTC::GFP (Fig. 6a-b). In control males, presence of female partners robustly induced nuclear CRTC::GFP accumulation in P1a neurons. By contrast, in *fru* mutant males, presence of females did not result in nuclear localization of CRTC::GFP in P1a neurons. Presence of male partners (either wild-type or *fru* mutant males) also did not cause CRTC::GFP nuclear localization. *in vivo* calcium imaging experiments further confirmed that P1a neurons of *fru* mutants did not become activated even when the experimenter placed a female (or male) stimulus in contact with the subject mutant males (Fig. 6c-h and Extended Data Fig. 5). Thus, P1a neurons of *fru* mutant males, although capable of eliciting courtship behavior ^34^, fail to be excited by either females or males. Previous studies indicate that *fru* functions during development to sculpt male-specific features of courtship circuits by regulating cell survival and axon guidance ^43–45^. We suggest that *fru* plays a role in establishing functional connections between sensory receptor neurons for female cues and P1a neurons, thereby linking detection of female cues to male arousal and courtship behavior. Male-directed courtship observed in *fru* mutants may be mediated by ectopic activation of neurons other than P1a, perhaps a different subset of P1 neurons ^34, 50^. Together, our results reveal that *fru* is necessary to ensure the activation of P1a neurons by female cues thereby specifying female-directed courtship, and highlight the utility of the CRTC::GFP reporter to efficiently examine neural correlates of behavior in wild-type as well as mutant flies.

**Fig. 6:**
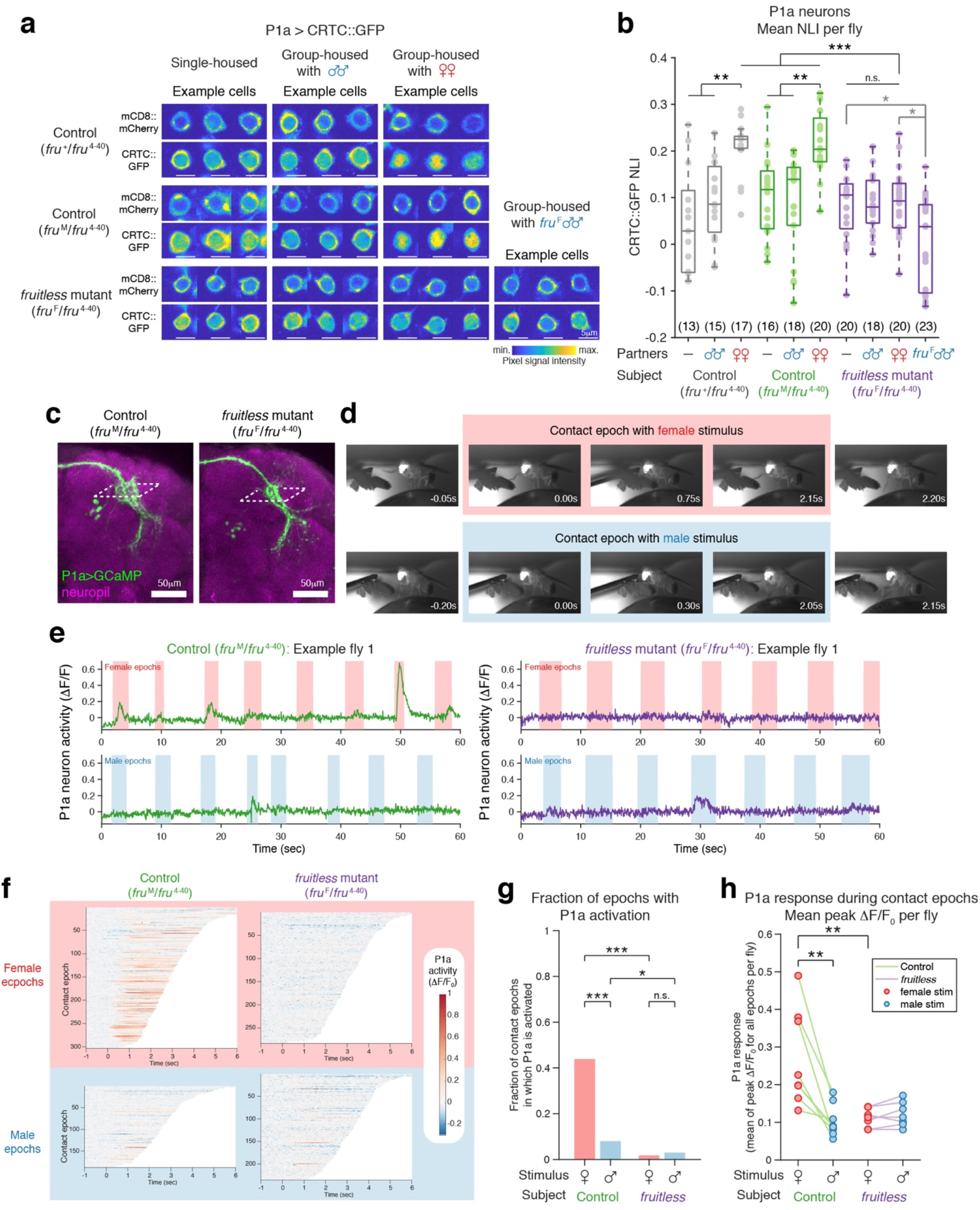
Examination of the activity of P1a neurons in *fruitless* mutants. (a) Representative confocal slice images of P1a neuron cell bodies in control vs *fru* mutant males in different social conditions. Flies were reared singly for 3 days upon eclosion and then paired with indicated partners for an additional 3-day period. (b) NLI of CRTC::GFP signal in P1a neurons of control vs *fru* mutant males in different conditions. Each dot represents mean NLI across cells per fly. Statistics by Kruskal-Wallis test followed by a post hoc Tukey’s HSD test; *** p<0.001, ** p<0.01, * p<0.05, n.s. p>0.05. (c) Confocal images of P1a neurons expressing GCaMP in control and *fru* mutant males. Dotted area represents approximate plane for calcium imaging. Anti-Brp antibody was used to label neuropil. (d) Example side view images of flies during calcium imaging experiments. Contact epoch (shaded pink and blue boxes) is defined as time between the first video frame in which a foreleg of the subject fly contacts the stimulus fly and the last video frame that the subject is in contact with the stimulus. (e) Example P1a activity upon contact with female (pink) or male (blue) abdomen in control (green) or *fru* mutant (purple) males. Shaded areas denote contact epochs. Median fluorescence intensity is used as baseline for calculation of ýF/F for this panel. See Extended Data Fig. 5a for additional example flies. (f) P1a activity in contact epochs. Time is relative to the initiation of contact with stimulus fly. Data are sorted by duration of contact epochs. Fluorescence intensity observed during one second prior to the contact initiation is used as the baseline for ýF/F0 calculation. See Extended Data Fig. 5b for unsorted data per fly. For control, 7 flies, 303 female epochs, 188 male epochs; for *fru* mutant, 7 flies, 283 female epochs, 235 male epochs. (g) Fraction of contact epochs in which P1a neurons become activated. Activation epoch if mean z-scored activity through epoch is over 1.96. See Extended Data Fig. 5c for data with a different threshold. Statistics by Fisher’s exact test; *** p<0.001, * p<0.05, n.s. p>0.05. (h) Mean P1a response across all contact epochs. Each dot is a fly and represents averaged peak ýF/F0 observed during each contact epoch for the fly. Statistics by Wilcoxon rank sum test between control and *fru* mutant and by Wilcoxon signed rank test between response to females vs males. ** p<0.01.

### Examination of neural activity in food-deprived flies using CRTC::GFP

Transcriptional neural activity reporters have been used to study neurons whose activity is regulated by internal states, such as hunger caused by food deprivation, and sleep drive caused by sleep deprivation ^10, 51, 52^. Our observation that sustained neuronal activation results in persistent nuclear localization of CRTC::GFP (Fig. 3a), whereas sustained neuronal silencing causes cytoplasmic localization (Fig. 3c), suggests that the CRTC::GFP reporter can also be used to examine neural correlates of long-lasting internal states. Because previous studies indicated that food deprivation induces dephosphorylation ^11^ and nuclear localization of CRTC ^53^, we wondered whether CRTC nuclear localization is a general starvation response observed across cells. We therefore examined the subcellular distribution of CRTC::GFP in P1a neurons of food-deprived flies (Fig. 7a). Food deprivation did not result in measurable increases in nuclear CRTC::GFP signals in P1a neurons (NLI for fed single-housed flies 0.16±0.06 vs food-deprived single-housed flies 0.17±0.05, p=1, Fig. 7b), whereas, as expected, housing with females led to increased nuclear localization as a result of neural activity (fed female-paired flies 0.29±0.04, p<0.001). These observations suggest that the subcellular distribution of CRTC::GFP provides a more sensitive readout of neural activity than starvation in P1a neurons. Curiously, female-dependent CRTC::GFP nuclear localization in P1a neurons was only observed in fed flies and not in food-deprived flies (NLI for female-paired food-deprived flies 0.20±0.05, p<0.05 compared to female-paired fed flies, Fig. 7b). This result indicates that P1a neurons in food-deprived males are not activated by the presence of females, suggesting that male courtship arousal is affected by feeding state, availability of food, or both (see ^54, 55^).

**Fig. 7:**
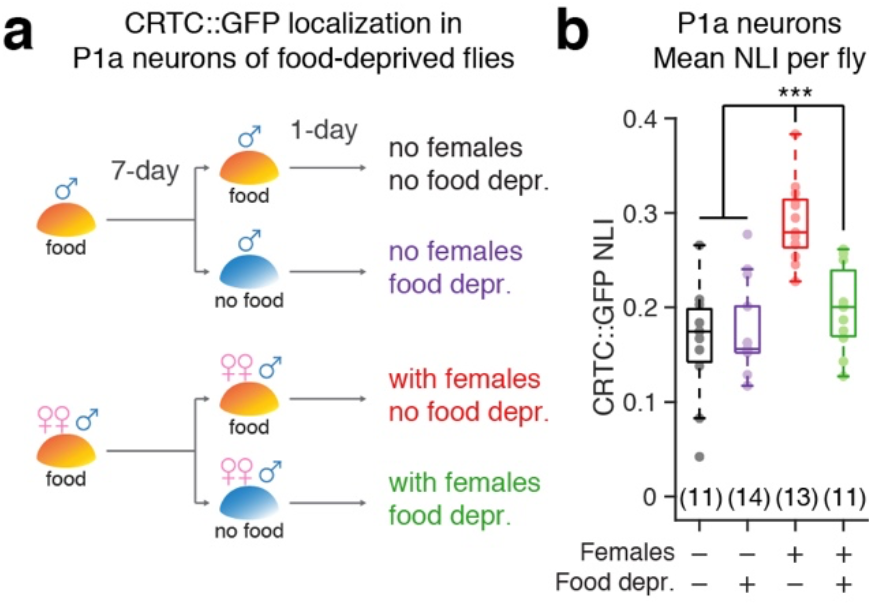
Examination of the activity of P1a neurons in food-deprived flies. (a) Schematic of experiment. Male subject flies were reared either singly or grouped with females for 7 days then food-deprived for one day. (b) Effect of food-deprivation and social conditions on subcellular localization of CRTC::GFP in P1a neurons. Statistics by Kruskal-Wallis test followed by a post hoc Tukey’s HSD test; *** p<0.001.

Together, our results suggest that, in addition to examining the neural correlates of behaviors, the CRTC::GFP reporter can be used to study neural activity modulated by persistent internal states including hunger.

## DISCUSSION

Activity-dependent gene expression promotes neural development as well as synaptic plasticity ^56, 57^. Membrane depolarization and resulting increases in local calcium levels activate cytoplasmic messengers, which transduce neuronal activation into transcriptional activation of multiple genetic loci. Transduction of these signals often depends upon translocation of molecular messengers over long distances. One clear example is mouse CRTC1 ^16, 17^. In silenced neurons, CRTC1 localizes to postsynaptic densities. Upon synaptic stimulation and resulting calcineurin activation, CRTC1 becomes dephosphorylated and is transported into the nucleus, where it regulates transcription of CREB target genes. Transcriptional activation, although rapid and called “immediate-early”, occurs over 30 minutes following neural activation ^4, 7^. Additional hours are necessary for the levels of transcripts or proteins to become detectable *in vivo* or *in situ*. In this study, we have exploited the transduction machinery that connects neural activation with transcriptional activation to develop a neural activity reporter that operates on a faster timescale of minutes.

Neural activity monitored by electrophysiological or optical recordings provides real-time changes in membrane potential or local calcium levels. These methods, however, require tethering and surgery, which often preclude reporting activity during natural behavior. Molecular reporters of neural activity provide useful alternatives to these recording techniques and allow for examination of neural activity that occurs during natural behavior. However, existing neural activity reporters depend on reporter gene expression. As a consequence, these reporters reveal neural activity that has occurred hours ago. Use of an early component in the transcriptional activation cascade that is not dependent on transcription or translation, but rather post-translational protein modification and translocation, provides an indicator of far more recent neural activity. Thus, the CRTC::GFP reporter permits measuring neural activity in behaving animals in a natural unrestrained environment.

The subcellular distribution pattern of CRTC::GFP persists when neural activity is persistent. Nuclear localization of CRTC::GFP persists for hours under conditions that continually activate neurons (Fig. 3a). In addition, we observed robust and sustained nuclear CRTC::GFP signals in P1a neurons of male flies housed with females for several days (e.g., Fig. 2f and Extended Data Fig. 2g). These results indicate that subcellular distribution of CRTC::GFP robustly reports persistent neural activation over a broad timeframe encompassing hours to days.

Different neurons exhibit different basal nuclear localization of CRTC::GFP that reflects tonic baseline activity. This suggests that the CRTC::GFP reporter can be used to detect decreased activity or inhibition. We indeed observed that upon removal of activating stimulus, CRTC::GFP rapidly redistributes from the nucleus to the cytoplasm (Fig. 3b) and that chronically silencing neurons results in cytoplasmic localization (Fig. 3c). Thus, the CRTC::GFP reporter is bidirectional and permits detection of both excitation and inhibition. The persistence and bidirectionality make CRTC::GFP an attractive reporter with which to examine neural correlates of longer lasting internal states.

It is noteworthy that we observed greater basal nuclear localization of CRTC::GFP in P1a neurons in older flies (different age flies were used for different experiments ranging from 3- days to 8-days old) (mean NLI in single-housed 3-day old males, -0.15 to -0.10 (Fig. 2e-f and Extended Data Fig. 2f-g), 6-day old males, 0.05 to 0.10 (Fig. 6a-b), 8-day old males, 0.16 (Fig. 7)). We do not know whether this is caused by an age-dependent increase in the basal neural activity of P1a neurons, changes in the expression and activity of molecules participating in CRTC signaling, or both. Nevertheless, in each of these experiments, we consistently observed that housing with females significantly increases nuclear CRTC::GFP signals, demonstrating the robustness of using the relative distribution of CRTC::GFP to detect neural activity.

Mouse CRTC1 and fly CRTC both translocate from the cytoplasm into the nucleus upon neuronal activity with similar kinetics ^16, 17^, raising the possibility that mammalian CRTC1 may also provide a tool for monitoring recent neural activity in mice and other mammals. More generally, activation of many signaling pathways induces translocation of molecules between the cytoplasm and the nucleus. These include activation of various developmental and neurotrophic signaling pathways ^58, 59^, steroid hormone receptors, such as glucocorticoid and estrogen receptors ^60–62^, and sterol regulatory element binding proteins that translocate from the endoplasmic reticulum to the nucleus upon sterol depletion ^63^. In many cases, nuclear translocation of effector molecules results in transcriptional changes and, as such, numerous cellular reporters have been developed using transcription as a readout (e.g., ^64, 65^). Given that nuclear import of effector molecules precedes transcription, subcellular-distribution-based reporters of signaling pathway activation afford faster response kinetics than transcription-based reporters. Establishing fly CRTC::GFP as a subcellular-distribution-based reporter of neural activity provides a model for developing reporters with which to monitor activation of different signaling pathways. Once developed, these reporters can be used for endpoint assays to compare signaling activity across animals in different conditions, as well as for *in vivo* imaging assays that monitor signaling activity within each animal during development or in response to environmental changes. Thus, future subcellular-distribution-based reporters may provide tools to study cellular-level activation of a broad spectrum of signaling pathways with more temporal precision than conventional transcription-based reporters.

Using CRTC::GFP, we have demonstrated activation of peptidergic neurons in male abdominal ganglia during mating, a function of the *fruitless* gene in ensuring activation of P1a neurons by the presence of females, and food-deprivation-dependent suppression of P1a activation. The automated segmentation of nucleus vs cytoplasm by a deep convolutional neural network was an essential analysis component that has facilitated quantification. While the present method requires individual cell body images as inputs, our future studies will focus on the development of an image processing pipeline that will allow for efficient quantification of CRTC::GFP from larger numbers of neurons using images with multiple cell bodies as inputs. A large-scale quantification pipeline, together with the bidirectionality of the CRTC::GFP reporter, may afford the ability to identify all neurons that are activated or inhibited during different behaviors. Thus, we envision that the CRTC::GFP reporter may help map the global landscape of cellular-resolution neural activity in the entire nervous system.

## ACKNOWLEDGEMENTS

We thank Woj Wojtowicz, Helmut Kramer, and Larry Zipursky for their comments on the manuscript; Camille Rogine and Rose Larios for technical assistance; Kenta Asahina, Yoshinori Aso, Gerry Rubin, Zhiguo Shang, Matthew Sieber, Dean Smith, and members of the Axel and Hattori laboratories for discussions; Yoshinori Aso for sharing fly stock prior to publication; David Anderson, Kenta Asahina, Wes Grueber, Marc Montminy, Gary Struhl, Bloomington Drosophila Stock Center and Vienna Drosophila Resource Center for fly stocks; UT Southwestern Bioinformatics Core Facility for help with machine learning, and Ron Doris, Clayton Eccard, Benjamin Fields, Miriam Gutierrez, Phyllis Kisloff, Adriana Nemes, and Leah Taylor for administrative assistance. This work is supported by UT Southwestern Endowed Scholarship to D.H. R.A. is a Howard Hughes Investigator.

## AUTHOR CONTRIBUTIONS

K.C.M., R.A., and D.H. conceived the project. M.B., K.J.S., M.G.M., A.Z., A.M., K.E.C., J.G.B., X.W., and D.H. performed experiments and analyzed data. A.R.J. and D.H. performed DCNN training. R.A. and D.H. supervised the project. D.H. wrote the manuscript with input from all authors.

## DECLARATION OF INTERESTS

The authors declare no competing interests.

## METHODS

### REAGENTS AND RESOURCES

| REAGENT or RESOURCE | SOURCE | IDENTIFIER |
| --- | --- | --- |
| Antibodies |  |  |
| Chicken-anti-GFP | Aves Labs | GFP-1020 |
| Rabbit-anti-DsRed | Clontech | 632496 |
| mAb anti-Bruchpilot | DSHB | nc82 |
| mAb anti-β-Galactosidase | Promega | Z3781 |
| mAb anti-Calceineurin (β-subunit) | Sigma | C0581 |
| AlexaFluor488 goat-anti-chicken | Life Technologies | A11039 |
| AlexaFluor568 goat-anti-rabbit | Life Technologies | A11036 |
| AlexaFluor633 goat-anti-mouse | Life Technologies | A21052 |
| Chemicals, peptides, and recombinant proteins |  |  |
| Farnesol | Sigma-Aldrich | F203 |
| Mineral Oil | Fisher Scientific | O122 |
| VectaShield | Vector Laboratories | H-1000 |
| SlowFade Gold | Invitrogen | S36938 |
| SlowFade Diamond | Invitrogen | S36963 |
| Experimental models: Organisms/strains |  |  |
| *D. melanogaster* : MB247-DsRed, UAS-GCaMP6f | <sup>66</sup> |  |
| *D. melanogaster* : UAS-CRTC::GFP in attP40 | This study |  |
| *D. melanogaster* : UAS-CRTC::GFP in attP40 | This study |  |
| *D. melanogaster* : UAS-GFP in attP40 | This study |  |
| *D. melanogaster* : UAS-mCD8::mCherry (2nd Chr.) | Gift from W. Grueber |  |
| *D. melanogaster* : UAS-nls::LacZ (2nd Chr.) | Gift from G. Struhl |  |
| *D. melanogaster* : UAS-eGFP::TORC | Gift from M. Montminy<br><sup>15</sup> |  |
| *D. melanogaster* : UAS-mCD8::mCherry-T2A-nls::LacZ in JK22C | This study |  |
| *D. melanogaster* : UAS-mCD8::mCherry-T2A-nls::LacZ in VK00005 | This study |  |
| *D. melanogaster* : UAS-dTRPA1 in attP2 | <sup>23</sup> |  |
| *D. melanogaster* : UAS-Kir2.1 (no tag) in VK00005 | This study |  |
| *D. melanogaster* : UAS-EGFP::Kir2.1 (3rd Chr.) | <sup>38</sup> |  |
| *D. melanogaster* : UAS-CanB/B2-shRNA in attP2 | TRiP at Harvard Medical School (BDSC)<br>67<br>68 | JF02616 |
| *D. melanogaster* : UAS-EGFP-shRNA in attP40 | TRiP at Harvard Medical School (BDSC)<br>67<br>68 | Bloomington 41552 |
| *D. melanogaster* : UAS-GCaMP6f in su(Hw)attP5 | This study<br>66 |  |
| *D. melanogaster* : SS01165 (R44A02-p65ADZp in attP40; VT033006-ZpG4DBD in attP2) | Gift from Y. Aso<br>22 |  |
| *D. melanogaster* : MB027B (R24H08-p65ADZp in attP40; R53F03-ZpG4DBD in attP2) | 24<br>66 |  |
| *D. melanogaster* : MB312B (R58E02-p65ADZp in attP40; R10G03-ZpG4DBD in attP2) | 24 |  |
| *D. melanogaster* : P1<sup>a</sup>-spGAL4 (R15A01-p65ADZp in attP40; R71G01-ZpG4DBD in attP2) | Gift from D. Anderson<br>29<br>30 |  |
| *D. melanogaster* : VT037580-GAL4 in attP2 | VDRC<br>66 |  |
| *D. melanogaster* : c155 (elav-GAL4) | BDSC<br>69 |  |
| *D. melanogaster* : Crz-GAL4 | Gift from D. Anderson<br>39 |  |
| *D. melanogaster* : P1<sup>a</sup>-spGAL4, *fru*<sup>4-40</sup> (R15A01-p65ADZp in attP40; R71G01-ZpG4DBD in attP2, *fru*<sup>4-40</sup>) | Gift from K. Asahina<br>34 |  |
| *D. melanogaster* : *Fru*<sup>M</sup> | Gift from K. Asahina<br>48 |  |
| *D. melanogaster* : *fru*<sup>F</sup> | Gift from K. Asahina<br>48 |  |
| Recombinant DNA |  |  |
| pUAST>stop>mCD8-GFP | Addgene | 24385 |
| pJFRC-MUH | Addgene | 26213 |
| pJFRC7-20xUAS-IVS-mCD8::GFP | Addgene | 26220 |
| 10xUAS-CRTC::GFP (in pJFRC-MUH) | This study |  |
| 10xUAS-GFP (in pJFRC-MUH) | This study |  |
| 10xUAS-mCD8::mCherry-T2A-nls::LacZ (in pJFRC-MUH) | This study |  |
| 20xUAS-Kir2.1 (no tag) (in pJFRC7 backbone) | This study |  |
| Software and algorithms |  |  |
| MATLAB | Mathworks |  |
| Fiji (ImageJ) | NIH |  |
| Subpixel Registration Algorithm | Mathworks File Exchange<br>70 |  |
| UbwonkoNet (ResNet-50 trained for subcellular pixel classification) | This study |  |

### RESOURCE AVAILABILITY

Data and data analysis codes including neural network will be deposited to GitHub. Further information and requests for resources and reagents should be directed to and will be fulfilled by the corresponding author, D.H.

### EXPERIMENTAL MODEL DETAILS

#### Flies

Driver lines MB027B, MB312B ^24^, VT037580-GAL4 ^66^, P1^a^-spGAL4 ^30^, SS01165 ^22^, c155 ^69^ and crz-GAL4 ^39^ were described previously. Effector lines UAS-GCaMP6f ^66^, UAS-dTRPA1 ^23^, and UAS-EGFP::Kir2.1 ^38^ were described previously. UAS-EGFP-shRNA and UAS-CanB/B2-shRNA (JF02616) were obtained from Bloomington Drosophila Stock Center (TRiP collection, ^67, 68^). *fruitless* mutant flies *fru*^M^, *fru*^F^ (both ^48^), and *fru*^4–40^ ^71^ were previously described.

#### Generation of new transgenic flies

10xUAS-CRTC::GFP. CRTC coding sequence and GFP sequence were PCR amplified and subcloned from genomic DNA of UAS-eGFP-TORC flies (gift from M. Montminy, ^15^) and pUAST>stop>mCD8::GFP ^72^, respectively. CRTC::GFP fusion sequence was generated by overlap extension PCR and these sequences were subsequently cloned into pJFRC-MUH vector ^73^ using NotI and XhoI. 10xUAS-GFP was generated by cloning the above GFP sequence into pJFRC-MUH vector using NotI and XhoI. UAS-CRTC::GFP and UAS-GFP were inserted in attP40 landing site using phiC31-mediated integration ^74^. 10xUAS-mCD8::mCherry- T2A-nls::LacZ. mCD8::mCherry and nls::LacZ sequences were PCR amplified and subcloned from genomic DNA of UAS-mCD8::mCherry flies (gift from W. Grueber) and UAS-nls::LacZ flies (gift from G. Struhl), respectively. These sequences were then PCR amplified with primers that contain a T2A self-cleaving (or ribosome-skipping) peptide sequence ^75^ and resulting PCR products were cloned into pJFRC-MUH at NotI site using Gibson assembly. UAS- mCD8::mCherry-T2A-nls::LacZ was inserted in VK00005 ^76^ and JK22C ^77^ landing sites.

20xUAS-Kir2.1 (untagged). Kir2.1 sequence was PCR amplified and subcloned from genomic DNA of UAS-EGFP::Kir2.1 flies ^38^, and this sequence was subsequently cloned into NotI site of modified pJFRC7 (20xUAS-IVS-mCD8::GFP) ^73^ without mCD8::GFP. UAS-Kir2.1 was inserted in VK00005 site. DNA sequences of all constructs were verified by Sanger DNA sequencing.

UAS-GCaMP6f in su(Hw)attP5 was generated by injecting the 20xUAS-GCaMP6f from ^66^ into su(Hw)attP5 landing site. All transgenic lines were generated by BestGene through phiC31- mediated integration.

### METHOD DETAILS

#### Calcium imaging

GCaMP imaging for DC3 PNs (Extended Data Fig. 1a) was performed as described in ^66^ using 2-photon microscope (Ultima, Bruker) equipped with ultra-fast laser (Chameleon Vision, Coherent). 3- to 5-day old female flies were used. Images were obtained with 60x/1.0NA water immersion objective (Olympus) with the following parameters: laser wavelength 925nm, laser power 2-4mW after objective, pixel size 0.39 microns, pixel dwell time 2 microsec, and frame rate 15.4fps. Odors were delivered using an olfactometer as described in ^66^. Farnesol was diluted in mineral oil at 1:10.

GCaMP imaging for P1a neurons was performed upon flies that were head-fixed and on an air-supported ball (10mm diameter) using 2-photon microscope (Investigator, Bruker) equipped with ultra-fast laser (Chameleon Ultra 2, Coherent). Males of the experimental genotype were isolated within one day upon eclosion, were reared singly, and were imaged when 2-3 day old.

Images were obtained with 20x/1.00NA water immersion objective (Olympus) with the following parameters: laser wavelength 925nm, laser power 10-15mW after objective, pixel size 0.13 microns, pixel dwell time 4 microsec, and frame rate 18.0fps. The fly-on-ball setup was made based on ^78^ with modifications. Detailed description of the fly-on-ball setup will be reported elsewhere. A male subject fly was ice-anaesthetized and head-fixed by the eyes using UV-cured glue (Bondic) to a metal shim (A-Laser) that was affixed to a custom 3D-printed saline reservoir. Proboscis was fixed in retracted position by UV glue. Head cuticle was removed in oxygenated saline (103mM NaCl, 3mM KCl, 5mM TES, 26mM NaHCO3, 1mM NaH2PO4, 1.5mM CaCl2, 4mM MgCl2, 8mM trehalose, 10mM glucose, adjusted to 270-275mOsm bubbled with 95%O2/5%CO2, pH7.2-7.3) using 27Gx1/2 hypodermic needle and trachea and fat over the brain were removed using forceps. Oxygenated saline was perfused over the brain continuously during imaging at approximately 0.5mL/min. Muscle 16 (frontal pulsatile organ) was pinched to minimize brain movement. Stimulus flies were virgin females or males (2U, w^1118^isoCJ1) that were 3- to 10-day old. Their legs and wings were removed, and they were glued by the thorax to a pin in a trifurcated stimulus holder mounted on a manipulator (Sutter). Two orthogonally placed cameras (front view and sideview of subject fly, Flea3, FLIR) were used to coordinate presentation of abdomen stimuli, and contact epochs between the stimulus and the subject fly were determined by post hoc frame-by-frame annotation of sideview video obtained at 20 frames-per-second using a custom MATLAB app (epoch durations; 3.1±1.6 sec, N=1009).

Imaging time was aligned to videorecording time by the onset and offset of 2-photon laser captured in the videos.

#### Live imaging of CRTC::GFP

2-photon live imaging of CRTC::GFP signals in DC3 PNs (Fig. 1b-e and Extended Data Fig. 1b) was performed using the set up described above for calcium imaging of DC3 PNs. 3- to 5-day old female flies were used. In each odor exposure block, 1 sec pulse of farnesol was presented 20 times with 5 sec interstimulus intervals. CRTC::GFP and mCD8::mCherry signals in the DC3 cell bodies were imaged before odor exposures (baseline) and after each odor exposure block. Images were acquired as z-stack with 1 micron interval and each slice was scanned 4 times to generate an averaged image with the following parameters: pixel size 0.13 or 0.19 microns, pixel dwell time 4 microsec. Each block was approximately 10 min including the imaging time.

Therefore, for control no odor condition, DC3 cell bodies were imaged every 10 min. NLI was calculated for each cell, and the difference between the NLI at each timepoint and at the baseline is represented as ýNLI.

#### Immunostaining and confocal microscopy

Experimental flies were cold anesthetized by transferring them into a pre-chilled vial and were dissected in 1xPBS. For experiments involved in determining CRTC::GFP subcellular localization, anesthesia and dissection were performed fly-by-fly to minimize differences in time between anesthesia and fixation. Fixation and immunostaining were performed as described in ^66^ using the following antibodies; chicken anti-GFP (1:1000, Aves labs), rabbit anti-DsRed (1:1000, Clontech), mAb anti-Bruchpilot (nc82, 1:10, DSHB), mAb anti-β-Galactosidase (1:1000, Promega), mAb anti-Calcineurin (ý-subunit) (1:100, Sigma) AlexaFluor488 goat-anti-chicken (1:200, Life Technologies), AlexaFluor568 goat-anti-rabbit (1:200, Life Technologies), and AlexaFluor633 goat-anti-mouse (1:200, Life Technologies). Immunostaining was performed to maximize the signal-to-noise ratio of CRTC::GFP and mCD8::mCherry and to avoid bleaching observed when native fluorescence of GFP or mCherry is examined. Samples were mounted in VectaShield (Vector Labs), SlowFade Gold, or SlowFade Diamond (Life Technologies) and were imaged using LSM510, LSM800, or LSM880 system (Zeiss). For calcineurin RNAi experiments examining the efficacy of knockdown (Extended Data Fig. 2b), control (GFP shRNA) vs experimental (CanB/B2 shRNA) brains were immunostained in the same wells to minimize staining variability and brains in the same well were distinguished by removing an optic lobe for one genotype. Images of cell bodies used for quantification of CRTC::GFP signals were all acquired as 12-bit images of 1 micron interval z-stack slices using a Plan-APOCHROMAT 63x/1.4 objective (Zeiss) with pixel size between 0.07 to 0.13 microns. For each z-stack, the imaging setting was adjusted using the brightest cell body, such that the pixel intensities of each channel spanned the entire dynamic range while avoiding oversaturation particularly for GFP channel. The lookup table for pseudo-colored images in figures was each adjusted for the minimum and maximum intensity.

#### Thermogenetic activation experiments with dTRPA1, silencing experiments with Kir2.1, and RNAi experiments

For dTRPA1 activation experiments, flies were reared at 21°C and genotyped one day before the experiments, when a single female fly of the experimental genotype is paired with a male in a fresh food vial. The experimental flies were 2- to 5-day old on the day of experiments. Flies were incubated either at room temperature (22-25°C) or in a water bath heated to 32°C for different durations before dissection and fixation. For nuclear export kinetics experiments, flies were incubated at 32°C for 15 minutes and then transferred to and incubated in vials at room temperature for different durations. For Kir2.1 silencing (Fig. 3c), 3- to 5-day old flies reared at 25°C and genotyped at least one day before the experiments were used. For RNAi experiments (Extended Data Fig. 2b-c), 3- to 7-day old flies reared at 25°C and genotyped within one day after eclosion were used.

#### P1a CRTC::GFP experiments

For experiments shown in Fig. 2e-f and Extended Data Fig. 2f, experimental males were genotyped within a day after eclosion and were housed either singly, with 2 males, or with 2 virgin females for 3 days (immunostained when 3- to 4-day old). For experiments shown in Extended Data Fig. 2g, experimental males were genotyped within a day after eclosion, were housed singly for 2 days, and were then transferred to and incubated in a new vial that contains either no fly, 2 male flies, or 2 virgin female flies for 1 day (immunostained when 3- to 4-day old). For experiments shown in Fig. 7, experimental males were genotyped within a day after eclosion, were housed either singly or grouped with 6 virgin females for 6 to 7 days, then were transferred (together with females for grouped condition) into vials that contain 1% agar in water (i.e., food-deprivation) and were housed for 18 hours before dissection, fixation, and immunostaining (as 7- to 8-day old). Control groups were transferred into vials that contain fresh food. For experiments shown in Fig. 6a-b, experimental males were genotyped within a day after eclosion, were housed singly for 3 days, and were then transferred to and incubated in a new vial that contains either no fly, 6 male 2U flies, or 6 virgin female 2U flies for 3 days (immunostained when 6- to 7-day old). For the condition in which multiple *fruitless* mutant males were cohoused, one or two males of the experimental genotype were housed with sibling *fruitless* mutant males in a group of 5-7 for 6 days.

#### Training convolutional neural networks for automated pixel classification

16 different GAL4 or spGAL4 driver lines representing a variety of cell types were crossed with UAS-CRTC::GFP; UAS-mCD8::mCherry-T2A-nls::LacZ flies to obtain experimental flies (performed as a part of screen, which will be reported elsewhere). A total of 774 brains (48.4±18.2 brains per line) were dissected, immunostained, and imaged using confocal microscope as described above. An experimenter determined boundaries for plasma and nuclear membranes using mCD8::mCherry and nls::LacZ signals with a custom MATLAB app for a total of 9127 cell bodies (570±487 cells per line), and these boundaries were used to assign each pixel a label as “outside”, “cytoplasm”, or “nucleus”. This app also saved experimenter-defined rectangle images that each captured a cell body whose boundaries were traced. These images were padded to adjust the image size such that it is appropriate as input for different convolutional neural networks. Images of these 9127 cell bodies excluding signals from the CRTC::GFP channel (i.e., only signals from mCD8::mCherry and nls::LacZ constituted each image) and corresponding labels were the input to train and evaluate convolutional neural networks for pixel classification. 8680 images, which contain randomly drawn 95% of cells for each driver line, were used for training, while the remaining 447 images were held out from training for performance evaluation. Training of networks was done with GPU using trainNetwork function of MATLAB Deep Learning Toolbox using the stochastic gradient descent with momentum (SGDM) optimizer with the following parameters; initial learning rate 10^-3^, mini batch size 144, number of epochs 3. Networks trained are ResNet-50 ^79^, ResNet-18 ^79^, MobileNetV2 ^80^, Xception ^81^, and U-Net ^82^. We evaluated the performance of each trained network with two metrics; confusion matrix to examine pixel-level label accuracy and correlation between NLIs calculated based on manually-annotated vs network-annotated ROIs (see below). Based on overall performance, we chose trained ResNet-50 (which we call “UbwonkoNet”) for automated pixel classification.

#### Corazonin CRTC::GFP mating experiments

Males of the experimental genotype were isolated upon eclosion and 2- to 8-day old males were used for experiments. Half of the males were paired with 2 virgin 2U females in normal food vial, while the other half were transferred into new food vial. Mating was scored manually (copulation occurred upon pairing an experimental virgin male with a virgin female within 15.6±12.6 min, N=16), and immediately upon copulation termination, males were ice anaesthetized and thoracic segments were dissected in 1x PBS and fixed for one hour. After this initial fixation, ventral nerve cord was further dissected and fixed for an additional 30 minutes. Cellular and nuclear boundaries were drawn manually using mCD8::mCherry signals. We observed significantly higher NLI in mated males regardless of their age, therefore the data are combined across age.

#### Copulation behavioral experiments

Males of the experimental genotypes were isolated upon eclosion and 4 to 7-day old males were used for experiments. 10- to 14-day old 2U virgin females (group-housed) were used as targets. Behavior experiments were performed in a custom-built 9-arena chamber comprising a 3D-printed frame and glass slides (Extended Data Fig. 4a). Female flies were anaesthetized by CO2 and one female per arena was loaded. Once females recover from anesthesia, males were loaded into each arena without anesthesia using a small 3D-printed funnel. Chamber was tapped continually while loading males to prevent mating. Recording was performed for 120 minutes using a camera (Flea3, FLIR) by taking one snapshot of the arenas every two seconds, except for 3 fly pairs for which one snapshot was taken every five seconds.

### QUANTIFICATION AND STATISTICAL ANALYSIS

#### Calcium imaging data analysis

Calcium imaging data were analyzed using custom MATLAB apps and scripts. Time-series images were first registered to a baseline averaged image using a subpixel registration algorithm ^70^. Each ROI over a cell body (for DC3 PN imaging) or neuronal arbors (for P1a imaging) was manually drawn on an averaged image post registration and mean GCaMP pixel intensities of the ROI were calculated per frame. These raw fluorescence traces were converted to ýF/F0 traces using averaged fluorescence over 1-second immediately before odor onset for DC3 PN imaging or contact onset for P1a imaging as baseline (F0). The response magnitude was quantified as averaged ΔF/F0 over 1.5-second period starting from 0.5-second upon odor onset for DC3 PN imaging or as peak ΔF/F0 during contact epoch for P1a imaging. For P1a imaging, we observed trial-by-trial variability in the likelihood and timing of activation in addition to response magnitude (see Fig. 6e and Extended Data Fig. 5a). Therefore, we also analyzed fraction of contact epochs in which P1a neurons become activated using two different thresholding methods. In both methods, the fluorescence trace for each contact epoch was converted to a z-scored trace using signals from 1-second immediately before contact epoch as baseline. In the first (Fig. 6g), a contact epoch was classified as activated if the mean z-score over the entire contact epoch is 1.96 or more. In the second (Extended Data Fig. 5c), a contact epoch was classified as activated if the minimum z-score of a sliding window of 5 consecutive frames (approximately 0.28-second) is 1.96 or more at any point during contact epoch (i.e., the fluorescence signal rose above 1.96x SD of baseline period for at least 0.28-second). Non-parametric tests were used for statistics (Wilcoxon rank sum test for unpaired data, Wilcoxon signed rank test for paired data, and Fisher’s exact test for categorical data. See Supplementary Table 2).

#### CRTC::GFP data analysis

Image data for CRTC::GFP experiments were analyzed using Fiji for experiments involving dTRPA1 or custom MATLAB apps and scripts for the other experiments. For manual drawing of boundaries that delineate plasma vs nuclear membranes, the analysis app only showed mCD8::mCherry (and nls::LacZ, where applicable) signal to minimize bias in ROI determination. For each cell body, experimenters chose a slice that best represents cross section of the cell body that contains nucleus and cytoplasm in a given z-stack and drew along the boundaries demarcating plasma and nuclear membranes using drawpolygon function. Mean CRTC::GFP signal intensities in the nucleus vs the cytoplasm were then calculated from pixels labeled as nucleus vs cytoplasm based on these boundaries. For pixel classification performed by UbwonkoNet, we used a different custom MATLAB app that selects rectangular ROI over each cell body of a confocal stack, which then automatically segments the image into outside, cytoplasm, and nucleus pixels and extracts mean CRTC::GFP signals in each subcellular compartment. Nuclear localization index (NLI) was then calculated by the following formula; (mean nuclear CRTC::GFP signal – mean cytoplasmic CRTC::GFP signal) / (mean nuclear CRTC::GFP signal + mean cytoplasmic CRTC::GFP signal). The NLI scales from -1 to +1, where +1 indicates all CRTC::GFP signals are in the nucleus, -1 indicates all signals are in the cytoplasm, and 0 indicates signals are equal in the nucleus vs cytoplasm. We chose this metric over more conventionally-used ratio (nuclear signal / cytoplasmic signal) to avoid non-linearity of ratio. We note that the NLI tends to be more positive than one may expect based on CRTC::GFP signals of each image (for example, the NLI for a cell with slightly higher CRTC::GFP signals in the cytoplasm may not be negative). This is likely caused by discounting of the cytoplasmic signals because the number of pixels lining the boundary between cell and outside is always more than the number of pixels lining the boundary between nucleus and cytoplasm, and cell-outside boundaries may contain pixels with no CRTC::GFP signals (i.e., outside the cell) whereas nucleus-cytoplasm boundaries always contain CRTC::GFP signals.

NLIs from all cells were averaged per fly for statistical tests. Non-parametric tests were used for statistics (Wilcoxon rank sum test for unpaired data with two conditions, Kruskal-Wallis test followed by post hoc Tukey’s HSD test for data with more than two conditions. See Supplementary Table 2).

#### Estimation of fraction of CRTC::GFP molecules in the nucleus

We calculated the fraction of CRTC::GFP molecules in the nucleus vs cytoplasm with the assumption that mean CRTC::GFP signal intensities correspond to concentrations of CRTC::GFP molecules in each subcellular compartment. Nuclear radii relative to cell radii were estimated by calculating mean distances between the center of each nucleus to the edges of nuclear ROI vs cellular ROI. This analysis showed that relative nuclear radii for immunostained cell images are 0.71±0.09 and those of live 2-photon imaging cell images are 0.70±0.06. We then calculated nucleus vs cytoplasm volumetric ratios by cubing the radii (e.g., with relative nuclear radius of 0.70, nuclear volume : cytoplasmic volume = 1: 1.92). For Extended Data Fig. 1c, we determined the fraction of nuclear CRTC::GFP molecules with different relative nuclear radii ranging from 0.55 to 0.85 for a range of typically observed NLIs (-0.50 to 0.50).

#### Behavioral data analysis

To analyze mating behavioral data, Fiji was used to identify frames that capture the first unilateral wing extension, copulation initiation, and copulation termination. Fractions of successful mating was equivalent across the control and experimental groups (24/50 for Crz>GFP, 38/60 for Crz>CRTC::GFP, and 28/54 for Crz>EGFP::Kir2.1; statistics by Fisher’s exact test, p>0.1 for all pairwise comparisons). Fly pairs that did not copulate were excluded from analysis. Latency to copulation (Extended Data Fig. 4c) was determined as elapsed time from the beginning of recording to the first frame that the fly pair exhibits copulation. Copulation duration was determined as elapsed time from the first to the last frame for which the fly pair exhibits copulation. Some fly pairs, especially those that expressed EGFP::Kir2.1 in corazonin neurons, did not terminate copulation within the 120-minute recording period. Censoring was not performed, however, and copulation duration was determined as elapsed time from the first copulation frame to the end of recording for these fly pairs. Kruskal-Wallis test followed by post hoc Tukey’s HSD test was used to determine statistical significance.

#### Performance evaluation of convolutional neural networks

Performance of each trained convolutional neural network was evaluated at two levels. First, we analyzed the confusion matrix that describe the accuracy of performance at the level of pixels. This was done with the held-out dataset using evaluateSemanticSegmentation function of MATLAB Deep Learning Toolbox (other output metrics of this function are reported in Supplementary Table 3). Second, we analyzed the correlation between CRTC::GFP NLIs determined manually vs by trained networks. Linear regression was fitted using fitlm function of MATLAB Statistics and Machine Learning Toolbox and regression coefficient, intercept, and coefficient of determination were compared to evaluate performance of each trained network.

For a small fraction of cells (<1%, up to 3 cells out of 447 cells used for performance evaluation), the networks failed to detect any pixels that correspond to either nucleus or cytoplasm, which resulted in inability to calculate NLI for these cells. We report this fraction as P[fail] in Fig. 4b and Extended Data Fig. 3a. We found that these cells typically exhibit little signals, especially in nls::LacZ channel, and the size and location of the cell body appear different from other cells of the same class strongly labeled by the driver. These observations suggest that cells that networks fail to calculate may represent those exhibiting spurious expression of effectors that may not have been driven consistently by the driver. Based on the overall performance, we chose trained ResNet-50, which we call “UbwonkoNet”, to automatically segment cell body images.

---

**Extended Data Fig. 1:**
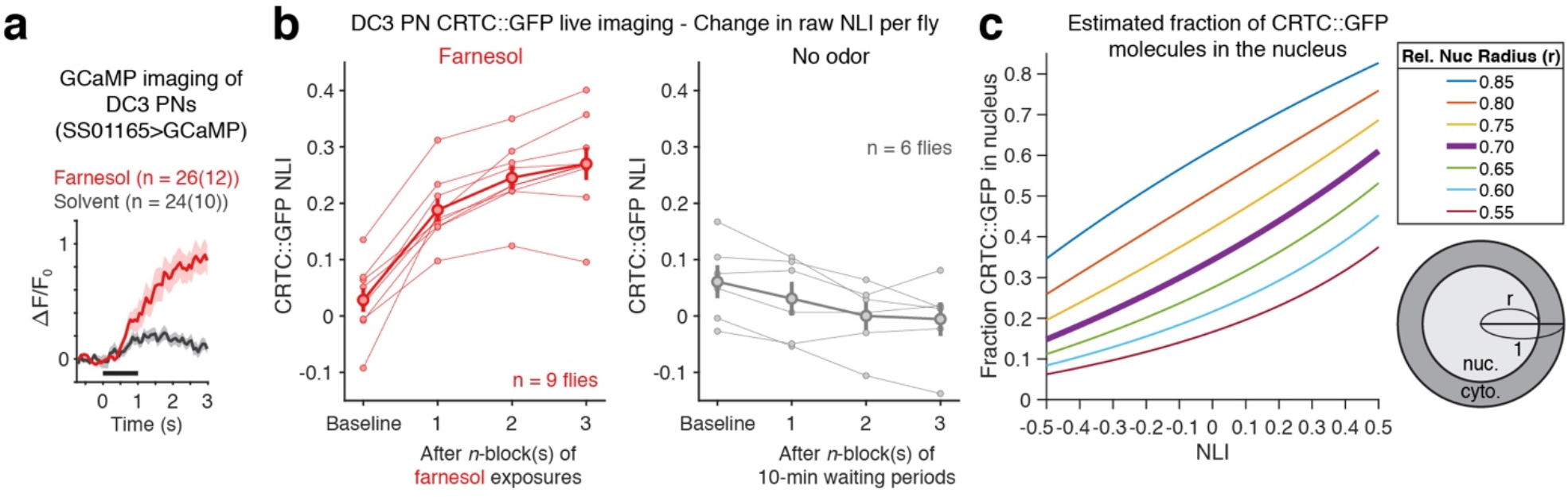
Translocation of CRTC::GFP in DC3 PNs. (a) ΔF/F0 traces from GCaMP calcium imaging experiments of DC3 projection neurons (PNs). Stimulus period is indicated by black line. Sample number represents the number of cell bodies imaged and number of flies in parenthesis. Lines indicate means and shadings indicate SEM. (b) Changes in CRTC::GFP nuclear signals (NLI) in DC3 PNs in different conditions. Thin lines represent individual flies (averaged across cells) and thick lines indicate mean ± SEM across flies. ΔNLI (Fig. 1d) was calculated for each cell by subtracting baseline NLI and averaged across cells per fly. (c) Estimated fraction of CRTC::GFP molecules in the nucleus for different NLIs in cells with different relative nuclear radii. The range of relative nuclear radii was determined to be 0.55 to 0.85 based on the cell and nuclear ROIs determined for 10,338 neurons of different cell types that were immunostained and imaged using confocal microscopy (0.71±0.09; mean ± SD). For DC3 live imaging, mean relative nuclear radius is calculated based on images obtained using 2- photon microscope (0.70±0.06; mean ± SD).

**Extended Data Fig. 2:**
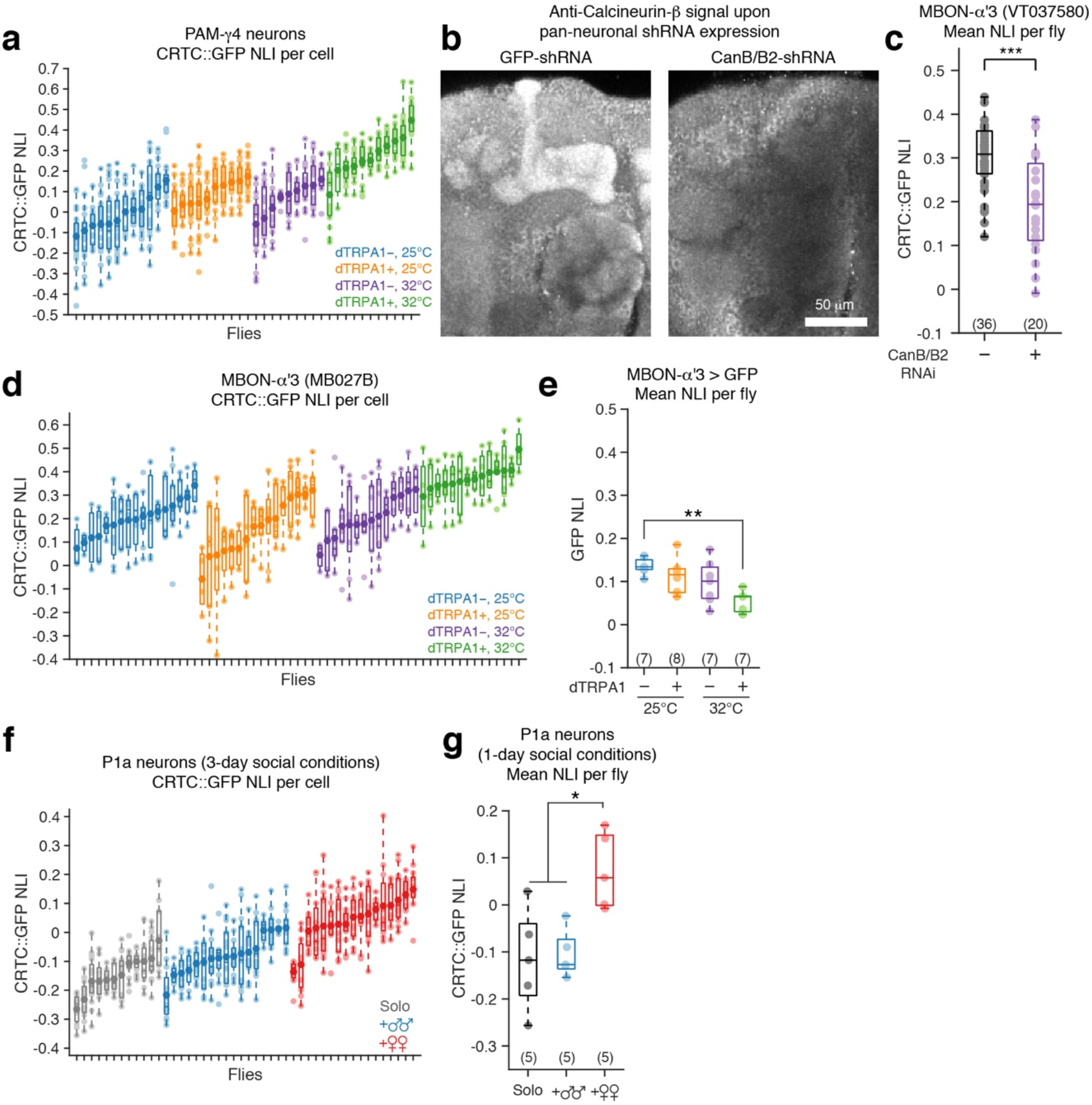
Subcellular distribution of CRTC::GFP provides a readout for neural activity in free moving flies. (a) NLI of CRTC::GFP signal in PAM-γ4 for thermogenetic activation experiments. Individual fly data for Fig. 2b. Lighter color dots are individual cells and darker color dots are mean of each fly. (b) Confocal images of brains immunostained with anti-calcineurin-ý antibody (see Methods). Maximum intensity projection images of slices that encompass antennal lobe to mushroom body lobes are shown. A pan-neuronal driver (c155, *elav*) was used to express shRNA targeting GFP (control, left) or CanB/B2 (right). Immunostaining for both control and experimental brains was performed in the same wells and the same imaging parameters were used for all brains. Representative images from 3 experiments (N=12 for GFP, N=12 for CanB/B2) are shown. (c) NLI of CRTC::GFP signal in the MBON-α’3 (labeled by VT037580-GAL4) in the presence of RNAi-mediated knockdown of calcineurin B and B2 (CanB/B2). Statistics by Wilcoxon rank sum test; *** p<0.001. (d) NLI of CRTC::GFP signal in MBON-α’3 for thermogenetic activation experiments. Individual fly data for Fig. 2d. (e) Thermogenetic activation experiments for MBON-α’3 using GFP instead of CRTC::GFP. Kruskal-Wallis test followed by a post hoc Tukey’s HSD test; p=0.002 for comparisons between dTRPA1+, 32°C and dTRPA1-, 25°C. (f) NLI of CRTC::GFP signal in P1a neurons of male flies in different rearing conditions. Individual fly data for Fig. 2f. (g) NLI of CRTC::GFP signal in P1a neurons of male flies in different rearing conditions. Subject flies were housed singly for 2 days upon eclosion and were housed for 1 day with males or females, or were left single-housed (“solo”). Statistics by Kruskal-Wallis test followed by a post hoc Tukey’s HSD test; * p<0.05.

**Extended Data Fig. 3:**
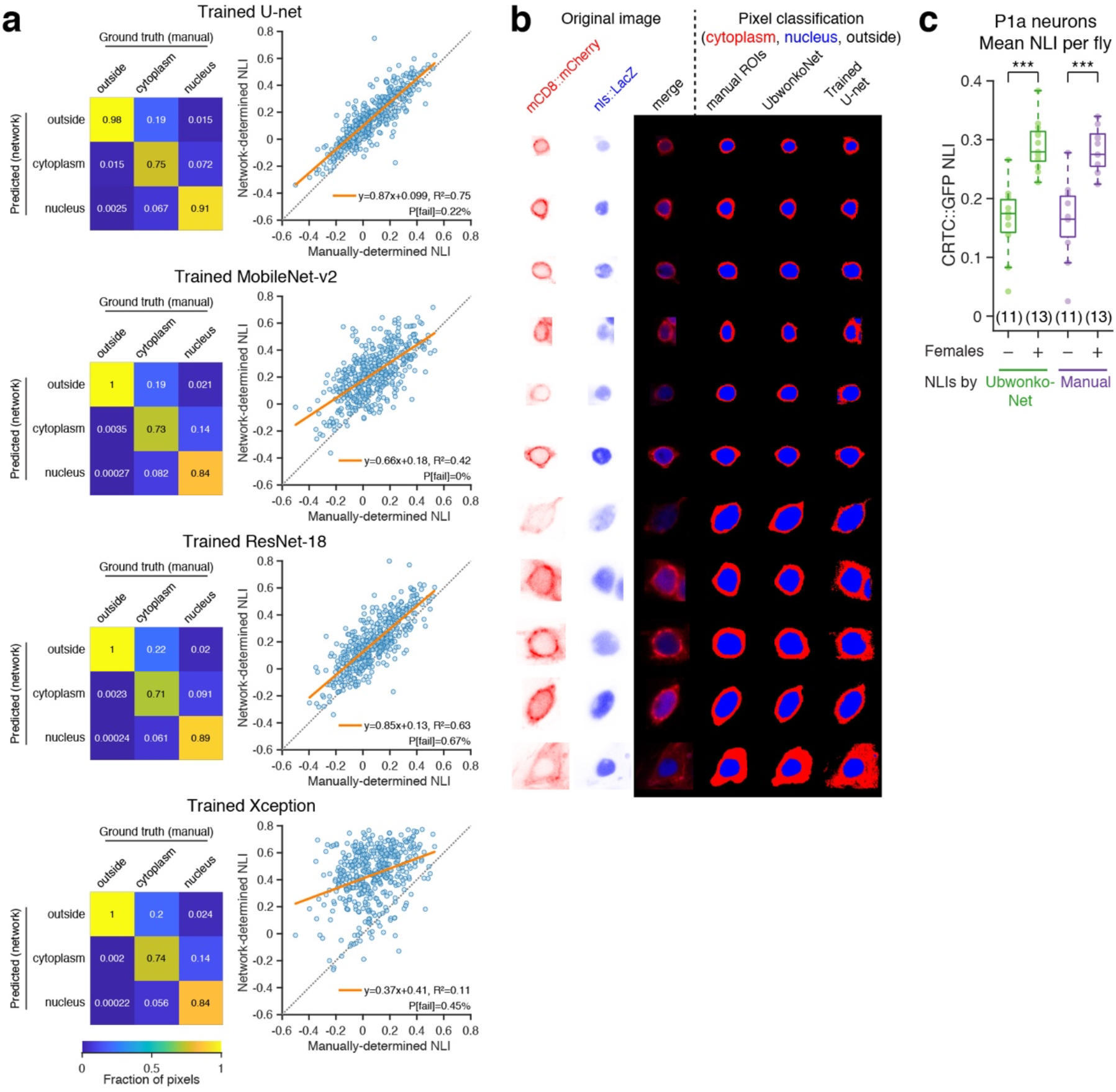
Evaluation of network performance. (Related to Fig. 4) (a) Performance evaluation for trained U-net, MobileNet-v2, ResNet-18, and Xception. (b) Eleven examples of pixel classification by trained ResNet-50 (UbwonkoNet) and trained U-net. The cells in the 4th and 8th rows show examples where U-Net failed to selectively “focus” on the cell body of interest. Examples include those shown in Fig. 4c. (c) Comparison of experimental results obtained using UbwonkoNet-determined NLIs vs manually-determined NLIs. A subset of data from the P1a experiments described in Fig. 7 (*ad libitum* fed group) were manually quantified (purple) and compared to the results obtained by UbwonkoNet (green). The left two columns are the same as the first and the third column of Fig. 7b. Statistics by Kruskal-Wallis test followed by a post hoc Tukey’s HSD test; *** p<0.001.

**Extended Data Fig. 4:**
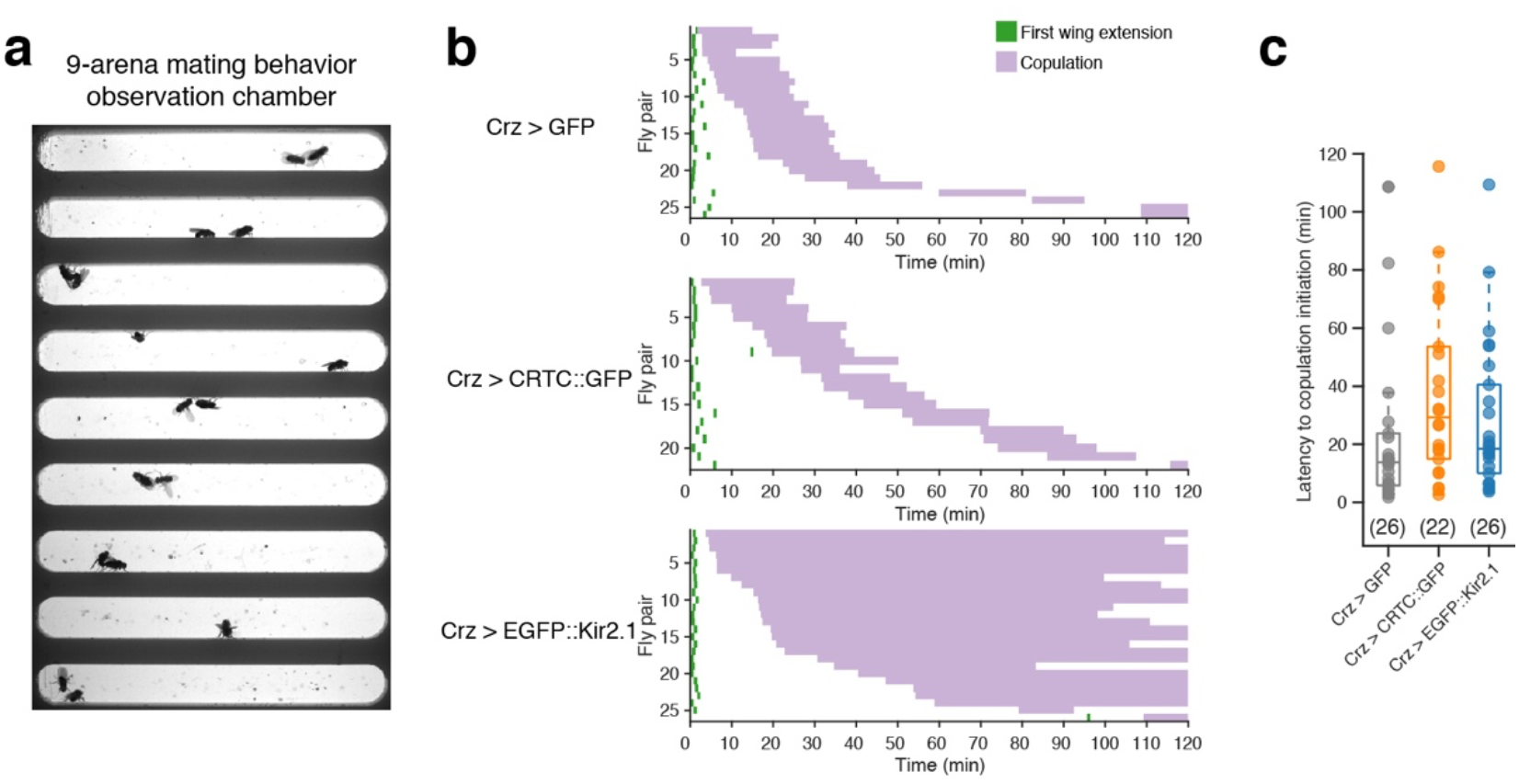
Expression of CRTC::GFP in corazonin neurons does not significantly affect mating behavior. (a) Behavior observation chamber used to examine mating behaviors. (b) Ethograms of mating experiments described in the right panel of Fig. 5b. Males of indicated genotypes were paired with wild-type females in the behavior observation chamber. (c) Latency to copulation initiation. Statistics by Kruskal-Wallis test followed by a post hoc Tukey’s HSD test; p>0.05.

**Extended Data Fig. 5:**
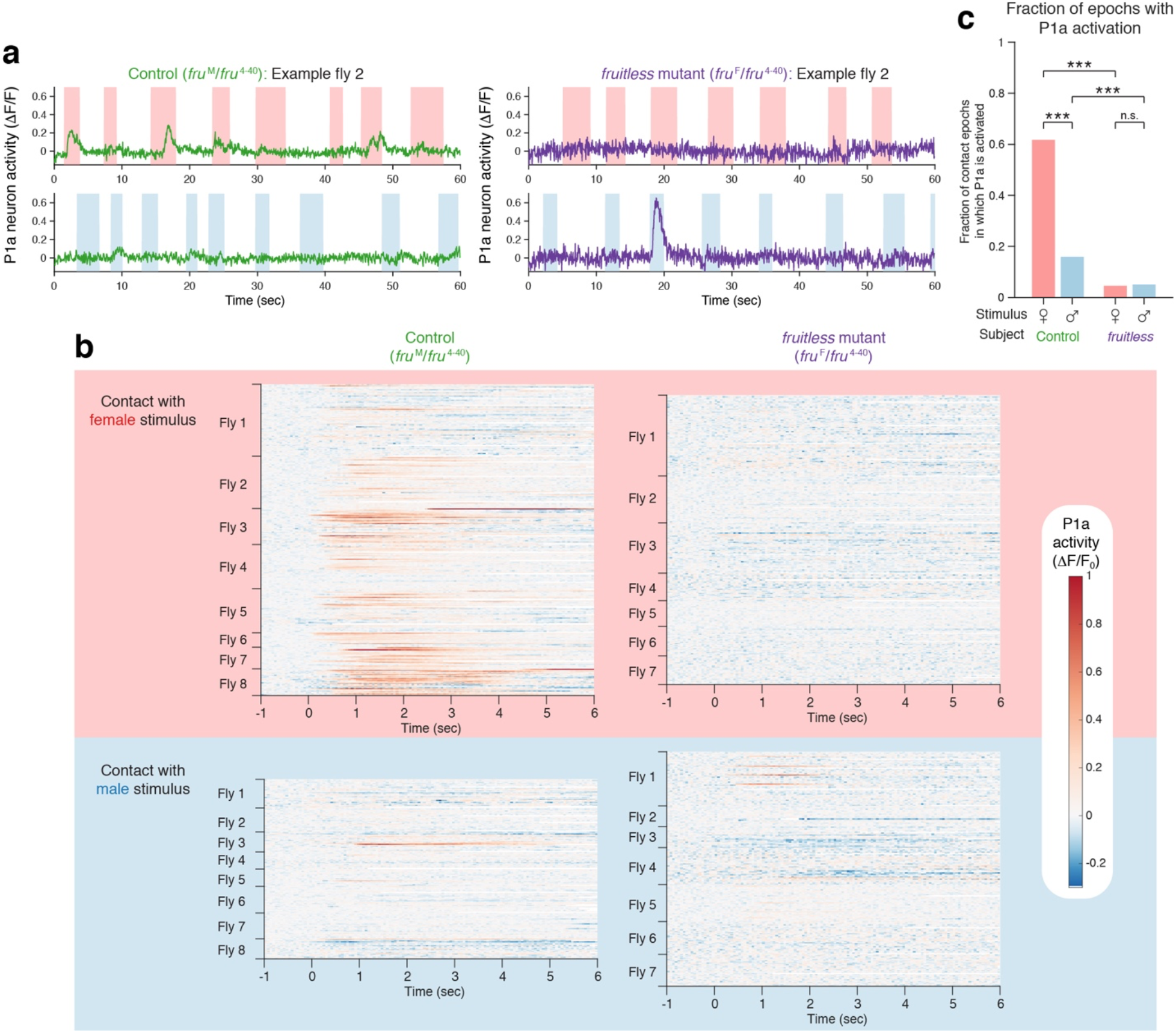
Examination of P1a activity in *fruitless* mutant flies. (a) Additional example flies for P1a calcium imaging. (b) P1a GCaMP response in contact epochs. Unsorted data for Fig. 6f. Calcium traces are shown only for the duration of each contact epoch, which varies from epoch to epoch. (c) Fraction of contact epochs in which P1a neurons become activated as determined with a different threshold from Fig. 6g. Activation epoch if at least 5 consecutive frames (appx. 0.28 sec) exhibit over 1.96 z-scored activity. Statistics by Fisher’s exact test. *** p<0.001, n.s. p>0.05.

## Notes

### Competing Interest Statement

The authors have declared no competing interest.

